# Two neuronal models of TDP-43 proteinopathy display reduced axonal translation, increased oxidative stress, and defective exocytosis

**DOI:** 10.1101/2023.05.17.540919

**Authors:** Alessandra Pisciottani, Laura Croci, Fabio Lauria, Chiara Marullo, Elisa Savino, Alessandro Ambrosi, Paola Podini, Marta Marchioretto, Filippo Casoni, Ottavio Cremona, Stefano Taverna, Angelo Quattrini, Jean-Michel Cioni, Gabriella Viero, Franca Codazzi, G. Giacomo Consalez

## Abstract

Amyotrophic lateral sclerosis (ALS) is a progressive, lethal neurodegenerative disease mostly affecting people around 50-60 years of age. TDP-43, a ubiquitously expressed RNA-binding protein involved in pre-mRNA splicing and controlling mRNA stability and translation, forms neuronal cytoplasmic inclusions in an overwhelming majority of ALS patients, of both sporadic and familial origin, a phenomenon referred to as TDP-43 proteinopathy. These cytoplasmic aggregates disrupt the subcellular transport and localization of mRNA. The axon, like dendrites, is a site of mRNA translation, permitting the local synthesis of selected proteins, both constitutively and in response to stimuli reaching the axon and presynaptic terminal. This is especially relevant in upper and lower motor neurons, whose axon spans long distances, likely accentuating their susceptibility to ALS-related noxae. In this work we have generated and characterized two models of TDP-43 proteinopathy, consisting of virtually pure populations of mouse cortical neurons expressing a human TDP-43 fusion protein, wt or mutant, which accumulates as cytoplasmic aggregates. Neurons expressing human TDP-43 exhibit a global impairment in axonal protein synthesis, an increase in oxidative stress, and defects in presynaptic function and electrical activity. These changes correlate with deregulation in the axonal levels of polysome-engaged mRNAs playing relevant roles in those processes. Our data support the emerging notion that deregulation of mRNA metabolism and of axonal mRNA transport may trigger the dying-back neuropathy that initiates motor neuron degeneration in ALS.

## 1. INTRODUCTION

Amyotrophic lateral sclerosis (ALS) is a progressive, lethal neurodegenerative disease affecting people around 50-60 years of age. The most common clinical signs are muscular weakness, spasticity, fasciculations and dysphagia, culminating in progressive paralysis and death (reviewed in Brown and Al-Chalabi, 2017; reviewed in Hardiman et al., 2017). The most frequent cause of death is respiratory failure (Gil et al., 2008; reviewed in Niedermeyer et al., 2019). About 10% of ALS patients also show cognitive impairment consistent with fronto-temporal dementia (FTD), which is characterized by degeneration of neurons of frontal and temporal lobes. More broadly, up to 50% of patients develop cognitive and/or behavioral impairments during disease progression e (reviewed in Brown and Al-Chalabi, 2017). Ten percent of ALS cases show familial inheritance (fALS), while the majority of patients, about 90%, have no family history and are classified as sporadic cases (sALS) (reviewed in Hardiman et al., 2017). ALS is characterized by the degeneration of upper (cortical) and lower (cranial or spinal) motor neurons.

Since 1994, many ALS causative genes and risk factors have been discovered. These genes can be grouped into three main categories based on the functional roles played by proteins involved in (1) protein homeostasis, (2) cytoskeletal maintenance, and (3) RNA metabolism. The involvement of RNA-binding proteins (RBPs) (Blair et al., 2010; Sreedharan et al., 2008) highlights the importance of altered RNA processing in ALS. TDP-43, a ubiquitously expressed RBP, has several functions: pre-mRNA splicing (Brown et al., 2022; Buratti et al., 2001; Fratta et al., 2018; Polymenidou et al., 2011; Tollervey et al., 2011; Watanabe et al., 2020), mRNA stability (Colombrita et al., 2012; Costessi et al., 2014; Strong et al., 2007), including that of its own transcript (Ayala et al., 2011), mRNA transport (Alami et al., 2014; Nagano et al., 2020), and the control of mRNA translation (Briese et al., 2020; Chu et al., 2019). TDP-43 is encoded by *TARDBP*, an essential gene (Kraemer et al., 2010) whose mutations account for about 5-10% of all forms of fALS (Sreedharan et al., 2008). However, while *TARDBP* mutations are relatively infrequent in ALS patients, TDP-43 is the main component of neuronal cytoplasmic inclusions found in 97% of all ALS patients, of both sporadic and familial origin. This phenomenon is referred to as TDP-43 proteinopathy (Arai et al., 2006; Neumann et al., 2006). TDP-43, a predominantly nuclear protein (reviewed in Cohen et al., 2011), preferentially binds UG-rich regions of RNA targets (Ayala et al., 2005; Buratti et al., 2001) and then shuttles to the cytoplasm (Ayala et al., 2008). Its C-terminal domain contains an intrinsically disordered protein region, the low complexity domain, which promotes liquid-liquid phase separation (LLPS) and leads to liquid droplet formation, a process that underlies the assembly of RNA granules (reviewed in Harrison and Shorter, 2017; Wolozin and Ivanov, 2019). Interestingly, the majority of ALS-mutations, located in the C-terminal domain (reviewed in Buratti, 2015; Prasad et al., 2019), affect the ability of TDP-43 to induce LLPS (Conicella et al., 2016) and may promote cytoplasmic aggregate formation, disrupting the subcellular transport and localization of its target mRNAs. While many proteins are synthesized in the neuronal soma and transported into the axon by fast or slow axonal transport (reviewed in Guo et al., 2020), the axon, like dendrites, is a site of local mRNA translation (reviewed in Jung et al., 2012; Martinez et al., 2019), and axonal and cell-body transcriptomes are compartment specific (Fusco et al., 2021; Shigeoka et al., 2019). Messenger RNAs are transported along the axon to synthesize selected proteins locally through the translation machinery (ribosomal subunits and whole ribosomes), whose composition undergoes remodeling in situ (Fusco et al., 2021; Shigeoka et al., 2019). Local translation occurs both constitutively and in response to stimuli acting far away from the cell body, in the axon and presynaptic terminal. Locally synthesized proteins include factors involved in axonal growth (Lee et al., 2018), axonal viability (Cosker et al., 2016), and translation (Fusco et al., 2021). While axonal translation was long considered a feature of developing or regenerating axons, it is now clear that it also plays a key regulatory and homeostatic role in mature axons (reviewed in Kim and Jung, 2020; Ostroff et al., 2019; Shigeoka et al., 2016). This is especially relevant in some upper and lower motor neurons, whose axon spans long distances, likely accentuating their susceptibility to stressful conditions and their absolute requirement for local response mechanisms largely independent of the cell body – indeed, in a meter-long axon, this compartment accounts for well over 99% of the total neuronal volume (reviewed in Ragagnin et al., 2019). Actually, several lines of evidence suggest that ALS is a distal axonopathy, characterized by axonal impairment, which precedes motor neuron degeneration and the onset of clinical signs (Fischer et al., 2004; Moloney et al., 2014).

The alteration of local mRNA translation in the axon may be a decisive factor in ALS progression, possibly affecting various cellular responses independently of the neuronal soma. For example, oxidative stress is a hallmark of ALS motor neurons, and an increase in reactive oxygen species (ROS) levels and ROS-associated damage have been reported in ALS (Altman et al., 2021; reviewed in Barber and Shaw, 2010; Colombrita et al., 2009; Lehmkuhl et al., 2021; Wong et al., 2021). While non-cell-autonomous mechanisms play important roles in the pathogenesis of this disorder or in its mitigation (reviewed in Beers et al., 2006; Liao et al., 2012; Liu and Wang, 2017; Marchetto et al., 2008; Wang et al., 2011), little is known about cell-autonomous signaling events controlling oxidative stress secondary to TDP-43 proteinopathy, and even less about the specific roles played by axons in the response to this condition.

Likewise, synaptic activity may be strictly dependent on local translation in the axon of long-range projection neurons. Indeed, it has been established that mature axons are enriched in transcripts involved in synaptic transmission: for instance, *UNC13A* (also known as *Munc13-1*) (Yang et al., 2015), a risk gene for ALS and FTD, participates in activity-dependent refilling of the readily-releasable vesicle pool and in neurotransmitter release, as demonstrated in mouse models (inter alia Augustin et al., 1999; Brown et al., 2022). Dysregulation of proteins involved in neurotransmitter release have been shown to precede denervation (Krishnamurthy and Pasinelli, 2021); hence, synaptic dysfunction may be a crucial factor in ALS onset and progression. Excitatory glutamatergic synapses have received special attention in the ALS field (Fogarty et al., 2016a; Fogarty et al., 2016b; Fogarty et al., 2017; Genc et al., 2016; Handley et al., 2017; Jiang et al., 2017); however, whereas numerous observations of early postsynaptic spine degeneration have been made, evidence is sparse regarding molecular and functional changes at the presynaptic terminal of corticospinal motor neurons.

Here, we describe a study conducted on cultured mouse cortical neurons expressing a human TDP-43 fusion protein, wt or mutant, which accumulates in the cytoplasm, forming insoluble granules. Neurons that accumulate TDP-43 in the form of insoluble cytoplasmic aggregates exhibit a global impairment in axonal protein synthesis, an increase in oxidative stress, and defects in presynaptic function and electrical activity, accompanied by deregulation in the axonal levels of polysome-engaged mRNAs playing relevant roles in those functional processes.

## 2. METHODS

### 2.1 Primary culture of cortical neurons

Animal handling and experimental procedures were performed in accordance with the EC guidelines (EC Council Directive 86/609 1987) and with the Italian legislation on animal experimentation (Decreto L.vo 116/92) and approved by our Institutional Animal Care and Use Committee.

Mouse embryonic cerebral cortices (strain C57BL/6N) were harvested at gestational day 14.5 (E14.5) and digested with Trypsin (Gibco, 15090-046). To obtain a single cell suspension the cells were mechanically dissociated with a glass Pasteur in neuronal culture medium (Neurobasal™ Plus Medium, Gibco A3582901, 1X B-27™ Plus Supplement, Gibco, A35828-01); the cell concentration was calculated with the T20 automated Cell Counter (Biorad). Cortical neurons were plated at the density of 42,000/cm^2^ on plastic or glass slides treated with Poly-D-Lysine (100μg/ml, Sigma, P6407), and cultured in an incubator at 37°C, 5% CO2. For specific experiments, cortical neurons were plated on microfluidic chambers (Xona Microfluidics, SND-450). In the cell body compartment, more neuron culture medium volume was added relative to the axonal compartment to generate hydrostatic pressure. At 2 days in vitro (DIV) 20μg/ml BDNF (Proteintech, 450-02-B) was added to the axonal compartment; fresh BDNF (10μg/ml) was added every three days.

### 2.2 Generation of lentiviruses expressing TDP-43 (wt and A315T)

For overexpression experiments in cortical neurons, we generated constructs encoding the wt human TDP-43 (hTDP-43) and the mutant human TDP-43 (A315T), fused N-terminally with the turbo-Red Fluorescent Protein (tRFP-hTDP-43 wt); the turbo-Red Fluorescent Protein (tRFP) was used as a control. For calcium assays control neurons were transduced with empty vector p277 instead of tRFP due to technical issues with the imaging instrument. The vectors were engineered in the laboratory, starting from the lentiviral vector pLenti277-GFP kindly provided by L. Naldini (San Raffaele-Telethon Institute of Gene Therapy, Milan, Italy). Lentiviral particles were prepared as described previously (Amendola et al., 2005). Briefly HEK293T cells were transiently co-transfected using the calcium-phosphate precipitation method with the transfer vectors, the MDLg/pRRE plasmid, the RSV-Rev plasmid and the MDLg plasmid encoding the G glycoprotein of the vesicular stomatitis virus. Cell supernatants containing lentiviral particles were collected 72 h after transfection, filtered and subjected to ultracentrifugation. The pellets were resuspended, divided into aliquots and stored at −80°C. To calculate the titer of lentiviral particles, cortical neurons were seeded in a 24-well plate and infected at 5 DIV with serial dilutions (10^-3^ to 10^-7^) of lentiviral particles overexpressing tRFP-TDP-43(wt), tRFP-TDP-43(A315T) and tRFP. Uninfected cortical neurons were used as controls. The experiment was repeated in triplicate for each condition. Cortical neurons were fixed after 9 days in vitro (DIV) and processed for immunofluorescence with tRFP and TuJ1 primary antibodies. ArrayScan microscope (Thermo Fisher ArrayScan XTI HCA Reader) was used to acquire 20 random fields/well. For every field we counted tRFP-positive infected cells (red) and the total number of cells, stained with a nuclear marker (DAPI). A ratio of total tRFP-positive cells to DAPI-positive cells was calculated for each well. To calculate the titer a given ratio of tRFP-positive cells to total DAPI positive cells between 0.1% and 10% (dynamic range) was chosen, and this formula was used: (Infected cells/Total cells) *Dilution factor *Number of seeded cells. In our experiments a multiplicity of infection (MOI) = 4 was used to achieve the transduction of most cells without detectable cellular toxicity. Cortical neurons were infected at 5 DIV and treated, fixed or lysed at different DIV based on the experimental needs.

### 2.3 Immunofluorescence (IF)

Cells were fixed with 4% Paraformaldehyde (PFA)/ 4% Sucrose in 1X PBS for 15 minutes, then incubated with primary antibodies in 10% Goat Serum (GS), 0.1% Triton in 1X PBS overnight at 4°C. For antibodies directed against nuclear epitopes a permeabilization step of 10 minutes with 0.5% Triton in 1X PBS was added. Subsequently, cells were incubated with secondary antibodies, washed three times with 1X PBS and counterstained with DAPI (D9542, Sigma). Images were acquired using Leica Confocal SP8 (Leica TCS SP8 SMD FLIM Laser canning Confocal), Nikon Spinning Disk (Nikon CSU-X1 Spinning Disk, Nikon TE2 inverted microscope), Axio Observer (Zeiss Axio Observer.Z1 with Hamamatsu EM9100). The quantification of all experiments was performed with the NIS-element Software (Nikon). When IF analyses were combined with heat shock treatments, cortical neurons were seeded in 24-well plates and infected as usual. At 14 DIV the plates were incubated in a water bath at 43°C for 60 minutes, washed in 1X D-PBS and processed for immunofluorescence.

### 2.4 Western blotting

Cortical neurons were scraped at room temperature using 50μl of Lysis Buffer (5% SDS, 10mM EDTA, 50mM Hepes pH 7.4) with the addition of protease inhibitors (Leupeptin, Sigma, L2884; Aprotinin, Sigma, A1153; PMSF, Sigma, P7626; Pepstatin A, Sigma, P5318; Sodium Orthovanadate, Sigma, S6508; Sodium Pyrophosphate, Sigma, 221368; Sodium Fluoride Sigma, S6776; β-Glycerophosphate Sigma, G6251). All protein extracts were sonicated and quantified with Pierce ™ BCA Protein Assay kit (ThermoFisher, 23225). 25μg protein lysate were resuspended in Laemmli buffer 1X (Sigma, S3401-1VL), denatured at 95°C for 5 minutes and loaded onto a polyacrylamide gel (Mini-PROTEAN(R) TGX ™ Precast Gels 4-20%, Biorad, 456-1094). Proteins were transferred to a nitrocellulose membrane (Biorad, 1704159) using Trans-Blot(R) Turbo™ (Biorad). Membranes were incubated in blocking buffer (5% milk in TTBS) for 1h at RT, then overnight at 4°C with primary antibodies diluted in blocking buffer. The membranes were incubated with secondary antibodies in blocking buffer. Bands were revealed using the Clarity ™ Western ECL Substrate (Biorad, 170-5061). The images were acquired using the ChemiDoc™MP (Biorad) and the quantification was performed with Image Lab ™Software (Biorad).

### 2.5 Primary and secondary antibodies

| Primary Antibodies for Immunofluorescence |  |  |  |  |
| --- | --- | --- | --- | --- |
| *Description* | *Brand* | *Cat. #* | *Host/ isotype* | *Dilution* |
| Map2 | Millipore | MAB3418 | Mouse/IgG | 1:1000 |
| TDP-43 | Proteintech | 10782-2-AP | Rabbit/IgG | 1:500 |
| βIII-Tubulin | Covance | MMS-435P | Mouse/IgG2a | 1:1000 |
| βIII-Tubulin | BioLegend | 802001 | Rabbit/IgG | 1:1000 |
| G3BP1 | Proteintech | 13057-2-AP | Rabbit/IgG | 1:300 |
| HuC/D | Thermo Fisher | A-21271 | Mouse/IgG2b | 1:2000 |
| tRFP | Evrogen | AB233 | Rabbit/IgG | 1:1000 |
| O4 | Sigma | O7139 | Mouse/IgM | 1:100 |
| CTIP2 | Abcam | ab18465 | Rat/IgG | 1:250 |
| GFAP | DakoCytomation | Z0334 | Rabbit/IgG | 1:500 |
| IBA1 | Wako | 01919741 | Rabbit/IgG | 1:500 |
| TBR1 | Abcam | ab31940 | Rabbit/IgG | 1:250 |
| GAD65 | GeneTex | GTX100281 | Rabbit/IgG | 1:100 |
| RPS6 | Cell signaling | mAb#2217 | Rabbit/IgG | 1:500 |
| RPL26 | BethylLab | A300-686AT | Rabbit/IgG | 1:500 |
| Caspase3 | BD PharMingen | 559565 | Rabbit/IgG | 1:200 |
| Puromycin | Millipore | MABE343 | Mouse/IgG | 1:100 |
| **Primary Antibodies for Western Blotting** |  |  |  |  |
| TDP-43 | Proteintech | 10782-2-AP | Rabbit | 1:1000 |
| Calnexin | Sigma | C4731 | Rabbit | 1:5000 |
| tRFP | Evrogen | AB233 | Rabbit | 1:5000 |
| GAPDH | Santa Cruz | sc-25778 | Rabbit | 1:2500 |
| TBR1 | Abcam | ab31940 | Rabbit/IgG | 1:1000 |

| **Secondary Antibodies for immunofluorescence** |  |  |  |
| --- | --- | --- | --- |
| *Description* | *Brand* | *Cat. #* | *Dilution* |
| Alexa Fluor™ 546 Goat anti-mouse (IgG) | Invitrogen | A11003 | 1:1000 |
| Alexa Fluor™ 546 Goat anti-mouse (IgG2a) | Invitrogen | A21133 | 1:1000 |
| Alexa Fluor™ 488 Goat anti-rabbit (IgG) | Invitrogen | A11034 | 1:1000 |
| Alexa Fluor™ 488 Goat anti-mouse (IgG) | Invitrogen | A11001 | 1:1000 |
| Alexa Fluor™ 488 Goat anti-mouse (IgG2a) | Invitrogen | A21131 | 1:1000 |
| Alexa Fluor™ 647 Goat anti-mouse (IgG) | Invitrogen | A32728 | 1:1000 |
| Alexa Fluor™ 647 Goat anti-rabbit (IgG) | Invitrogen | A21244 | 1:1000 |
| **Secondary Antibodies for Western Blotting** |  |  |  |
| Goat anti-mouse IgG (H+L)-HRP Conjugate | Biorad | 172-1011 | 1:10000 |
| Goat anti-rabbit IgG (H+L)-HRP Conjugate | Biorad | 170-6515 | 1:10000 |

### 2.6 Subcellular fractionation

All described procedures were performed at 4°C. Cortical neurons were washed in 1xPBS, scraped and lysed for 10 minutes in Lysis Buffer (50mM Tris–HCl pH 8.0, 10mM NaCl, 5mM MgCl2, 0.1% Nonidet P-40) with the addition of protease inhibitors. The lysate was spun at 1000 x g for 15 minutes to pellet the nuclear fraction (P1). The supernatant (S1) was centrifuged at 50.000 rpm for 60 minutes in a TLA55 rotor (Beckman) to yield crude cytosol (S2) and crude membrane pellet (P2). The pellet (P2) was resuspended in Lysis buffer without Nonidet P-40 and centrifuged again at 50.000 rpm for 60 minutes to yield washed crude membrane pellet (P2’). The nuclear fraction (P1) was washed three times with Lysis buffer without Nonidet P-40 at 1000 x g for 15 minutes. Pellet was resuspended in a high-salt buffer (20mM HEPES pH 7.5, 0.5M NaCl, 1mM EDTA, 1mM dithiothreitol and protease inhibitors) and rotated for 20 minutes. The pellet was centrifuged at 17.700 g for 30 minutes and the supernatant, containing the nuclear extract, was collected.

### 2.7 Puromycylation assay

DIV 14 cortical neurons plated on microfluidic chambers were treated with 2μM puromycin (Sigma, P8833) for 5 min. Pulse-labeling with a low concentration of puromycin, a structural analog of aminoacyl-tRNAs that is incorporated by nascent polypeptide chains, causing peptide release and disassembly of the two ribosomal subunits, was used as a readout of new protein synthesis (David et al., 2012). After treatment, the cultures were fixed and processed for immunofluorescence using an anti-puromycin primary antibody. Puromycin mean pixel intensity was measured in the soma, in 30 μm axon shaft and in the growth cone of at least 10 neurons per condition for each experiment.

### 2.8 Colocalization analysis

Colocalization analysis between tRFP-tagged TDP43 and endogenous HuC/D, RPS6 and RPL26 was performed in the soma of DIV 9 cortical neurons. A threshold was defined for each channel and the area colocalizing was normalized on the total TDP43 area. The analysis was performed on at least 10 neurons per condition for each experiment.

### 2.9 ROS measurement

Cortical neurons, plated in a 96-well plate (Greiner Bio-One, 655090) and cultured up to DIV 13, were loaded with 5μM dichlorofluorescein diacetate (CM-H2DCFDA, Thermofisher, C6827), a probe that turns into a fluorescent molecule (2’,7’-dichlorofluorescein, DCF) upon oxidation by intracellular ROS. The CM-H2DCFDA loading was performed in KRH pH 7.4 buffer (125mM NaCl, 25mM Hepes/NaOH pH 7.4, 5mM KCl, 1.2 mM MgCl2, 2mM CaCl2, 6mM Glucose) for 30 minutes at 37°C, followed by 10 minutes of incubation with the nuclear dye 1X Hoechst (Sigma, 33258) (Vannocci et al., 2018). For iron overload, neurons were treated with 20μM ferric ammonium citrate for 2 days before the analysis (Codazzi et al., 2016). The images were acquired using ArrayScan (ThermoFisher ArrayScan XTI HCA Reader) and the fluorescence intensity analysis was carried out by the ALEMBIC Facility of the San Raffaele Hospital.

### 2.10 Electrophysiology

Individual slides with cortical neuronal cultures were placed in a recording chamber mounted on the stage of an upright BX51WI microscope (Olympus) equipped with differential interference contrast optics (DIC). Cultures were perfused with artificial cerebrospinal fluid (ACSF) containing (in mM): 125 NaCl, 2.5 KCl, 1.25 NaH_2_PO_4_, 2 CaCl2, 25 NaHCO3, 1 MgCl_2_, and 11 D-glucose, saturated with 95% O_2_ and 5% CO_2_ (pH 7.3). ACSF was continuously flowing at a rate of 2-3 ml/min at 32°C. For mEPSC recordings, ACSF was added with the Na+ channel specific blocker tetrodotoxin (TTX, 1μM). Whole-cell patch-clamp recordings were performed in cortical neurons using pipettes filled with a solution containing the following (in mM): 124 KH_2_PO_4_, 10 NaCl, 10 HEPES, 0.5 EGTA, 2 MgCl_2_, 2 Na2-ATP, 0.02 Na-GTP, (pH 7.2, adjusted with KOH; tip resistance: 4-6 MΩ). All recordings were performed using a MultiClamp 700B amplifier interfaced with a computer through a Digidata 1440A (Molecular Devices, Sunnyvale, CA, USA). Traces were sampled at a frequency of 10 kHz and low pass filtered at 2 kHz. Data were acquired using Clampex software (Molecular Devices) and analyzed with Clampfit and GraphPad Prism applications. Statistical comparisons were obtained using SigmaStat 3.5 (Systat, San Jose, CA).

### 2.11 Calcium analyses

Cortical neurons were incubated with 4 μM fura-2 acetoxymethyl ester (AM, Calbiochem, CAS 108964-32-5) 40 min at 37°C; the ratiometric properties of fura-2 (excitation at 340nm and 380 nm and emission at 510 nm) permit the analysis of intracellular Ca2+ levels both at basal conditions and upon stimulation with 100 μM glutamate. Imaging setup and analysis are the same of FM1-43 assay. Single-cell video imaging setup consists of an Axioskope 2 microscope (Zeiss, Oberkochen, Germany) and a Polychrome IV (Till Photonics, GmbH, Martinsried, Germany) light source. Fluorescence images were collected by a cooled CCD video camera (PCO Computer Optics GmbH, Kelheim, Germany). The ‘Vision’ software (Till Photonics) was used to control the acquisition protocol and to perform data analysis (Codazzi et al., 2006; Rosato et al., 2022).

### 2.12 FM1-43 assay

Neurons were loaded with fura-2 AM as previously described, before being subjected to the FM1-43 assay. FM1-43 is a styryl dye whose loading and release from synaptic vesicles provides a reliable measurement of vesicle release. The neurons were treated with 20 μM FM1-43 (Sigma, cat. SCT126)(Betz and Bewick, 1992), dissolved in high K+ (60 mM)-containing KRH (HK-KRH; Na+ concentration was adjusted to maintain isotonicity), diluted 1:1 in normal KRH to obtain the required final KCl and FM1-43 concentration (30 mM and 10 μM respectively). FM1-43 was kept in the extracellular solution for 2 minutes, to complete endocytosis; this step was followed by several washes with dye-free KRH buffer, to eliminate the excess of FM1-43; after an additional stimulation with HK-KRH, fluorescence decay of FM1-43 over time was measured. The imaging setup is the same described for fura-2 calcium assay. FM1-43 fluorescence analyses were performed as described (Lazarenko et al., 2018).

### 2.13 Electron microscopy analysis

Neuronal cultures were fixed with 4% paraformaldehyde (PFA) and 2% (wt/vol.) glutaraldehyde in cacodylate buffer 0.12 mol/l pH 7.4 overnight at 4 °C, followed by incubation at room temperature for 2 h in 1% (wt/vol.) OsO4.,1,5 % potassium ferrocyanide in 100mM cacodylate buffer pH 7,4 for 1h on ice. After dehydration in a graded series of ethanol preparations, tissue samples were cleared in propylene oxide, embedded in epoxy medium (Epoxy Embedding Medium kit; Sigma-Aldrich, St. Louis, MO 63103 USA), and polymerized at 60 °C for 72 h. From each sample, one semi-thin (1 μm) section was cut with a Leica EM UC6 ultramicrotome (Leica Microsystems, Vienna, Austria). Ultra-thin (60 nm thick) sections of areas of interest were then obtained, counterstained with uranyl acetate and lead citrate, and examined with a transmission electron microscope (Talos 120C Fei), image were acquired with a 4kx4k Ceta CMOS camera (Thermo Fisher Scientific).

### 2.14 Transcriptome and translatome analysis

Miniaturized sucrose gradients were used to isolate free or polysomal RNA from cell bodies and axons of CNs cultured in microfluidic chambers, as described above. The procedure for polysome profiling by miniaturized sucrose gradient fractionation was adapted from published protocols (Negro et al., 2018). Sequencing data were retrieved from (Lauria, Maniscalco et al., in preparation) and processed as described by the authors. Briefly, reads were first preprocessed for adapter removal and trimming, then mapped to the mouse genome (GRCm38.p6, ENSEMBL release 92 and Gencode M17 gene annotation) and finally deduplicated. After sample size normalization based on the trimmed mean of M-values method), only genes with FPKM > 10 in all replicates of at least 1 sample were kept for subsequent analysis. Differentially expressed genes were identified by the edgeR applying multiple thresholds (CPM > 0.05, absolute log2 fold change > 0.75, p-value < 0.05). Annotation enrichment analysis with Gene Ontology terms, REACTOME and KEGG pathways were performed with the clusterProfiler Bioconductor package. GO analysis was performed as described (Bernabo et al., 2017).

Gene ontology tables were searched using the following query words: translation, ribosom* and polysom* for mRNA translation; oxidative, peroxid* and reactive oxygen species for oxidative stress; synap*; vesicle, exocyt*, endocyt* and neurotransmitter for synaptogenesis and synaptic function.

### 2.15 Statistical analysis

All values are expressed as mean ± standard error of the mean (SEM) of at least 3 independent experiments. Statistical analysis was performed using the GraphPad Prism software. The statistical tests for each experiment are reported in the figure legends. Non-significant differences yielding a p value ≥ 0.05 were regarded as non-significant. With respect to decay curves (Fig. 6), the model was fitted by restricted maximum likelihood. This model allows an easy interpretation of fluorescence decay because α_ctr, 〖α〗_ctr+Δ_(hTDP-43)) and〖α〗_ctr+Δ_(mutTDP-43)) are related to the fluorescence half-life in the three experimental conditions. Furthermore, multiplicative log-normal distributed errors address for the higher variability observed in correspondence of higher mean values.

## 3. RESULTS

### 3.1 E14.5 primary cortical cultures are strongly enriched in deep-layer glutamatergic neurons

To investigate the molecular mechanisms involved in the onset and progression of TDP-43 proteinopathy, we developed an *in vitro* model of highly homogeneous murine upper motor neurons (UMNs), which were lentivirally transduced to express human TDP-43, wild-type (wt) or carrying an ALS-mutation (Gitcho et al.), and N-terminally fused with turbo Red Fluorescent Protein (tRFP). To obtain primary cortical cultures enriched in glutamatergic neurons of layers V and VI, cells were isolated from the cerebral cortex of mouse embryos harvested at embryonic day 14.5 (E14.5), a developmental stage characterized *in vivo* by a low number of GABAergic neurons and glial cells in the dorsal forebrain. Indeed, the immunostaining performed at 3 DIV (**Fig. 1A**) revealed that virtually all cells were positive for the neuron-specific β3-tubulin (Ab: TuJ1), and 97% of them stained double-positive for the T-Box Brain Transcription Factor 1 (TBR1), an early marker of deep-layer glutamatergic neurons (Bedogni et al., 2010). Moreover, about 64% of neuronal cells were double-positive for TuJ1 and COUP-TF-interacting protein 2 (CTIP2), a transient marker of neurons located in cortical layers V and VI (reviewed in Molyneaux et al., 2007). To quantify the GABAergic component of these cultures, we immunostained these cells for glutamic acid decarboxylase-65 (GAD65). As shown in **Fig. 1B (**left panel), GAD65-positive cells were virtually absent, while they were abundant in neurons harvested at a later developmental stage (E17.5, **Fig. 1B**, right panel). Moreover, the results of EdU incorporation revealed that, while a small number of proliferating progenitors were still present at 2 DIV, all cells lost the ability to replicate DNA at 7 DIV (**Fig. 1C**).

**Figure 1.**
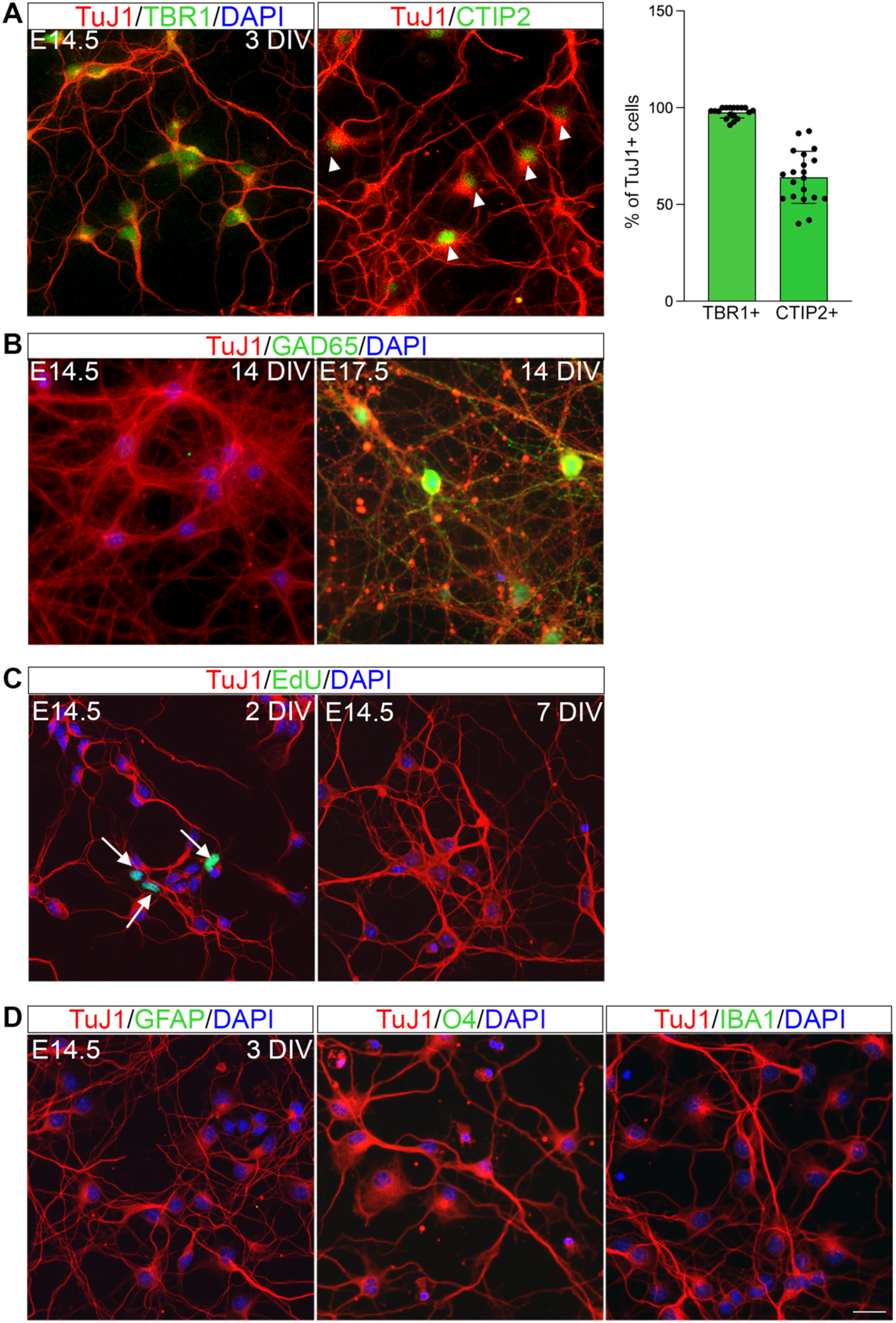
E14.5 primary cortical cultures are strongly enriched in deep-layer glutamatergic neurons. (A) Cortical neurons, established from E14.5 mouse embryos and cultured until 3 DIV, immunostained with a Tbr1 Ab (green) or with a CTIP2 Ab (green), and with the TuJ1 Ab (red), detecting β3-tubulin; blue: DAPI to visualize nuclei. The histogram on the right represents the percentage of TuJ1+ cells positive for TBR1 (97%) and CTIP2 (64%). Total number of neurons counted: 986 (18 fields) for Tbr1 and 912 (20 fields) for CTIP2 (data from 3 biological replicates). (B) left: UMN cultures harvested at E14.5, cultured for 14 DIV, fluorescently immunostained with the GAD65 Ab (green), a GABAergic neuron marker. No GAD65 signal is detected; right: cultures established from E17.5 mouse cortices are positive for GAD65 after the same time in vitro. (C) EdU staining of E14.5 UMN cultures after 2 DIV (on the left) and 7 DIV, to count proliferating cells. While some proliferating cells are present at 2 DIV (arrows), no EdU signal is detected after 7 DIV, indicating mitotic quiescence. (D) Representative images of UMN cultures at 3 DIV immunostained for GFAP (astrocytes), O4 (oligodendrocyte progenitors), or IBA1 (microglia) (green). No glial cells were detected in our cultures. Positive controls in Supplemental figure 1. Size bar: 25μm.

Finally, we evaluated the presence of glial cells, which are mostly produced at later stages of cortical development. TuJ1 immunostaining, combined with antibodies against specific glial markers (glial fibrillary acidic protein, GFAP, for astrocytes; oligodendrocyte marker 4, O4, for oligodendrocyte precursors; ionized calcium binding adaptor molecule 1, IBA1, for microglia) revealed the absence of all glial cell types in our mouse UMN cultures (**Fig. 1D**, positive controls in **Supplemental Fig. 1**).

In summary, E14.5 cortical neuronal cultures are strongly enriched in postmitotic glutamatergic neurons of deep cortical layers, from which the corticospinal tract arises, and contain a negligible contribution of glial cells or GABAergic neurons.

### 3.2 hTDP-43 and mutTDP-43 neurons give rise to cytoplasmic TDP-43 aggregates

A neuronal model of TDP-43 proteinopathy was produced by transducing the above UMN cultures with lentiviral particles to deliver tRFP alone (ctr neurons) or human tRFP-TDP-43, both wt and carrying the A315T mutation (henceforth defined hTDP-43 and mutTDP-43 neurons, respectively); constructs are sketched in **Fig. 2A**.

**Figure 2.**
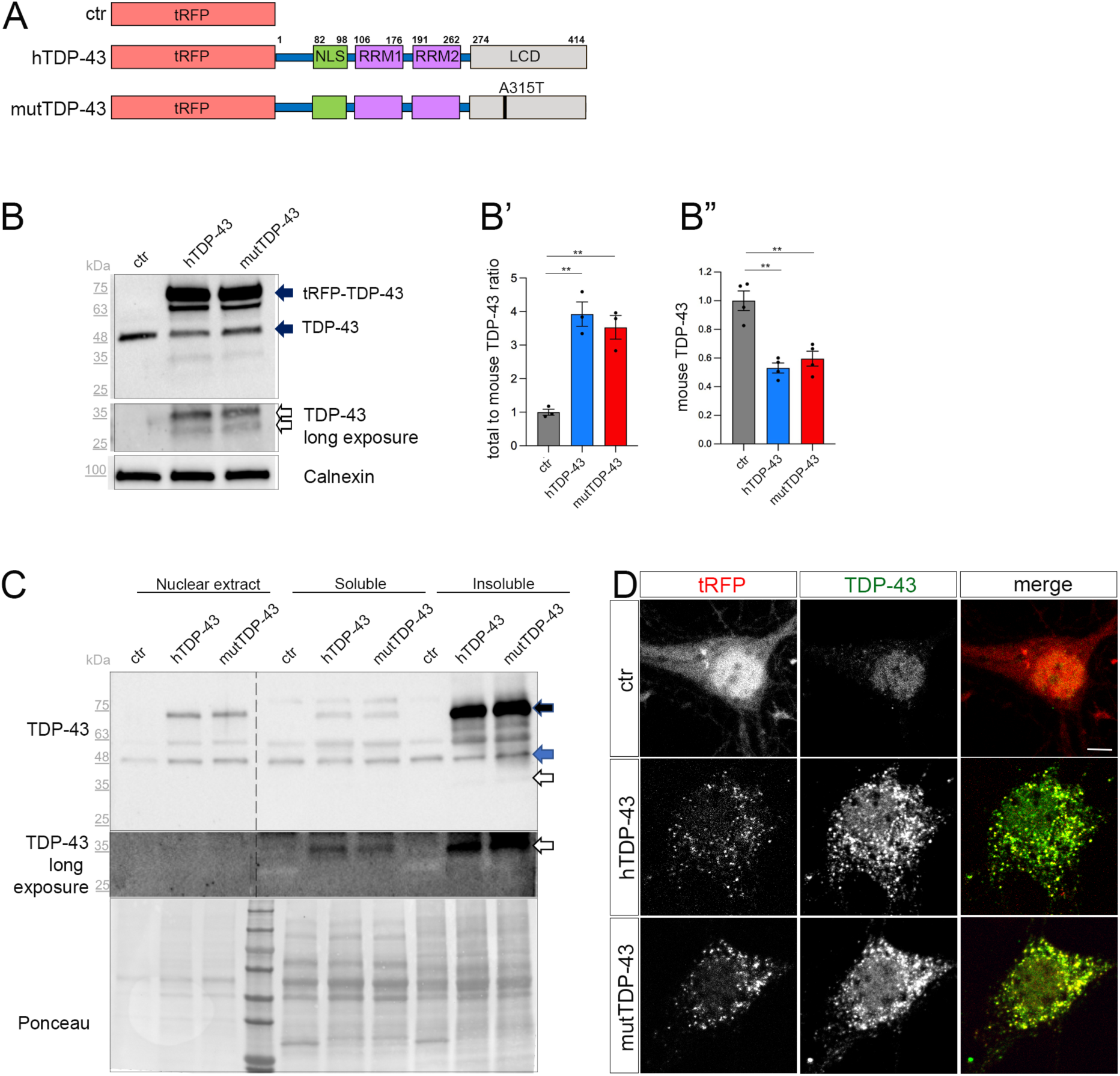
Cytoplasmic aggregate formation in neurons expressing tRFP-TDP-43. (A) Schematic representation of ctr, hTDP-43 and mutTDP-43 constructs. In hTDP-43 and mutTDP-43 constructs, TDP-43 functional domains are also shown. This schematic shows the presence of the fluorescent tag (tRFP) at the N-terminal position of TDP-43. NLS: Nuclear Localization Sequence; RRM1, RRM2: RNA Recognition Motif-1 and −2; LCD: low-complexity glycine-rich domain. The A315T aa substitution is located in the TDP-43 LCD domain. (B) Western blot of protein lysates derived from ctr, hTDP-43 and mutTDP-43 neurons immunostained with an Ab detecting both human and murine TDP-43. An anti-calnexin Ab was used as a loading control. The ∼75kDa band (upper solid arrow), corresponds to the fusion protein tRFP and human TDP-43 (wt or A315T), and a ∼48kDa band corresponding to endogenous murine TDP-43 (lower solid arrow). Another band (∼63kDa) is present in TDP-43-overexpressing neurons, likely representing a C-terminal cleavage product of the fusion protein. Upon a longer exposure, a ∼35kDa band (empty arrows) appears. (B’) The graph shows the ratio of total TDP-43 levels (fusion human+mouse endogenous) to endogenous TDP-43 levels in ctr cells; total TDP-43 signal in overexpressing neurons is about 3.5-3.8 times higher than in ctr ones. (B”) shows the normalized level of endogenous TDP-43 in TDP-43-overexpressing neurons relative to ctr ones; the endogenous protein level is halved relative to ctr cells, likely due to post-transcriptional autoregulation. Mean±SEM; B’: n=3; B”: n=4; One-way ANOVA test, **p<0.01. (C) Western blot of nuclear extract, cytoplasmic soluble fraction and insoluble fraction (see Methods) of ctr, hTDP-43 and mutTDP-43 neurons immunostained with a TDP-43 Ab. The exogenous protein (black solid arrow) is strongly enriched in the insoluble cytoplasmic fraction, while the endogenous protein (blue solid arrow) band is evenly present in all fractions. Likewise, the 30 kDa proteolytic fragment (empty arrow) is enriched in the insoluble cytoplasmic fraction of hTDP-43 and mutTDP-43 neurons. Ponceau staining is used as a loading control. (D) Ctr, hTDP-43 and mutTDP-43 neurons at 14 DIV, immunostained with an Ab detecting both human and murine TDP-43 (green). In ctr neurons, tRFP is concentrated mostly in the nucleus, while exogenous tRFP-TDP-43 (wt or A315T) is highly enriched in the cytoplasm of hTDP-43 and mutTDP-43 neurons, forming aggregates. Size bar: 5μm.

Since the cytoplasmic accumulation of TDP-43 is a key feature of ALS neurons (Suk and Rousseaux, 2020), we assessed the relative expression levels of the exogenous TDP-43 fusion proteins and investigated their subcellular localization. Biochemical analysis, performed on UMN lysates with an antibody recognizing both human and murine TDP-43, showed a marked overexpression of total TDP-43 protein (about 4-times higher than the endogenous protein in ctr cells; **Fig. 2B**, solid arrows, quantified in **Fig. 2B’**); notably, the level of endogenous murine TDP-43 is halved in hTDP-43 and mutTDP-43 neurons, probably due to negative post-transcriptional autoregulation (**Fig. 2B”**). A ∼65kDa band likely represents a degradation product of the fusion protein. Furthermore, a more prolonged exposure of chemiluminescence-stained membranes revealed, both in hTDP-43 and mutTDP-43 neurons, the presence of TDP-43 proteolytic fragments, a typical hallmark of ALS neurons (**Fig. 2B**, empty arrows) (Neumann et al., 2006).

To further characterize TDP-43 expression, we set up a subcellular fractionation protocol to separate nuclear from soluble and insoluble cytosolic fractions (**Supplemental Fig. 2**). The results of western blotting performed on subcellular lysates display an enrichment of exogenous hTDP-43 and mutTDP-43 proteins in the insoluble fraction (**Fig. 2C**, solid black arrow), together with their cleavage products (empty arrow), while endogenous murine TDP-43 is evenly present in all fractions analyzed (solid blue arrow). The cytoplasmic accumulation of exogenous TDP-43 was confirmed by immunocytochemistry, revealing a predominantly nuclear localization of the endogenous TDP-43 in ctr neurons (Neumann et al., 2006) and an enrichment of both hTDP-43 and mutTDP-43 in cytoplasmic aggregates (**Fig. 2D**).

Overall, our results indicate that neurons expressing RFP-TDP-43 fusion proteins recapitulate several key features of TDP-43 proteinopathy.

### 3.3 Exogenous TDP-43 is recruited to RNA granules

The analysis of tissue samples obtained from 97% of ALS patients has revealed the presence of TDP-43 pathologic inclusions (Ling et al., 2013; Neumann, 2009). TDP-43 is known to bind other RBPs (e.g., fragile X mental retardation protein, FMRP, and neuron-specific Elav-like Hu RNA-binding protein D, HuD), generating granules that transport mRNAs along neurites (Fallini et al., 2012). Therefore, we investigated the nature of exogenous TDP-43 aggregates in our UMN cultures by immunostaining hTDP-43 and mutTDP-43 neurons with an HuD- specific Ab. HuD is involved in mRNA stability, splicing, and positive regulation of translation (Diaz-Garcia et al., 2021). Our results reveal that exogenous TDP-43 granules partially colocalize with HuD (**Fig. 3A**). The degree of colocalization, plotted in **Fig. 3A’**, shows that HuD colocalizes slightly more frequently with hTDP-43 than with mutTDP-43.

**Figure 3.**
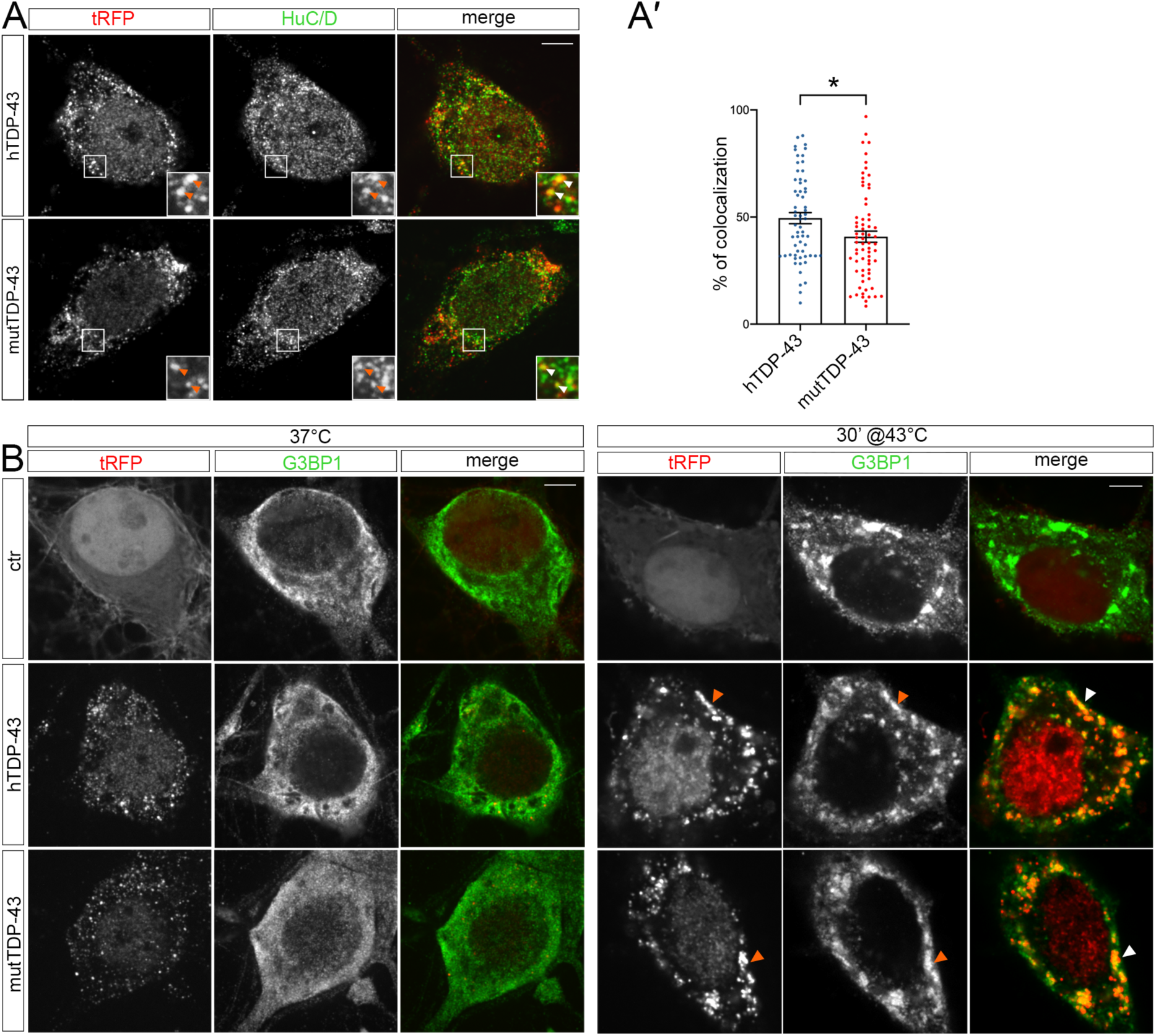
tRFP-TDP-43 is recruited to RNA granules. (A). Immunofluorescent staining of hTDP-43 and mutTDP-43 neurons with an anti-HuC/D antibody. HuC/D (green) partially colocalizes with fluorescent tRFP-TDP-43 aggregates (wt or A315T) (red) as shown by arrowheads in insets. (A’) dots in histogram indicate the percentage of HuC/D-tRFP colocalization in each individual cell examined. (B) Immunofluorescence of ctr, hTDP-43 and mutTDP-43 neurons with anti-G3BP1 under physiological conditions (left) and after heat shock (right). Under basal conditions G3BP1 (green) shows a diffuse pattern in hTDP-43 and mutTDP-43 neurons as in ctr cells. However, after heat shock (30’ at 43°C), G3BP1 generates stress granules that recruit exogenous TDP-43 (wt and A315T alike) (red) as shown by arrowheads. Size bar: 5μm. Mean ±SEM, n=3, Mann-Whitney test,*p-value<0.05.

To assess whether exogenous TDP-43-positive cytoplasmic granules colocalize with stress granules (reviewed in Dewey et al., 2012), we immunostained cells for Ras-GTPase-activating protein SH3 domain-binding protein 1 (G3BP1)), a known component of stress granules that is involved in their formation (Matsuki et al., 2013). Under basal conditions (37°C), G3BP1 signal displays a diffuse pattern in ctr, hTDP-43 and mutTDP-43 neurons alike, while heat shock treatment (43°C, 30 minutes) generated G3BP1+ granules that contained exogenous TDP-43 (**Fig. 3B**). This result recapitulates findings obtained in cells undergoing non-lethal injury (Higashi et al., 2013) and suggests that in our conditions TDP-43 does not *per se* promote stress granule formation but retains the ability to be recruited to newly assembled stress granules, once they form (Colombrita et al., 2009), although TDP-43 oligomerization and aggregation takes place in the cytoplasm separate from SGs (Streit et al., 2022).

### 3.4 Global downregulation of protein synthesis in the axon of TDP-43-overexpressing mouse cortical neurons

In parallel to the functional analysis of our cellular models (see below), we took advantage of a transcriptome and translatome analysis (Lauria, Maniscalco et al., in preparation) conducted on cell-body and axonal compartments, to explore the gene expression and mRNA translation landscape at a systems biology level. To this end, cortical neurons were plated on microfluidic chambers (Taylor et al., 2005), consisting of two main channels separated by 450μm-long microgrooves (see Methods); cell bodies are too large to enter them, while dendrites are too short to extend beyond them.

The physical separation of axonal and somatodendritic compartments was demonstrated by immunofluorescence in untransduced neurons, cultured in microfluidic chambers for 9 DIV and immunostained with Abs for Microtubule Associated Protein 2 (MAP2), a dendrite-specific marker, and for TuJ1, which decorates dendrites, cell bodies and axons alike. As shown in **Supplemental Fig. 3**, MAP2 signal is sharply restricted to the whole-cell channel, while axons extend into the right channel and are labeled by TuJ1 only.

To ask whether exogenous TDP-43 might sequester components of the mRNA translation machinery, possibly affecting local mRNA translation in the axon, we immunostained hTDP-43- and mutTDP-43-expressing neurons with antibodies against small and large ribosomal subunit proteins (RPS6 and RPL26, respectively). As shown in **Fig. 4A,A’**, 20% of granules, on average, colocalize with RPS6 and RPL26, in both hTDP-43 and mutTDP-43 neurons. This finding suggests that TDP-43 aggregates may sequester ribosome components and/or interfere with ribosomal transport, likely resulting in an overall decrease of mRNA translation.

**Figure 4.**
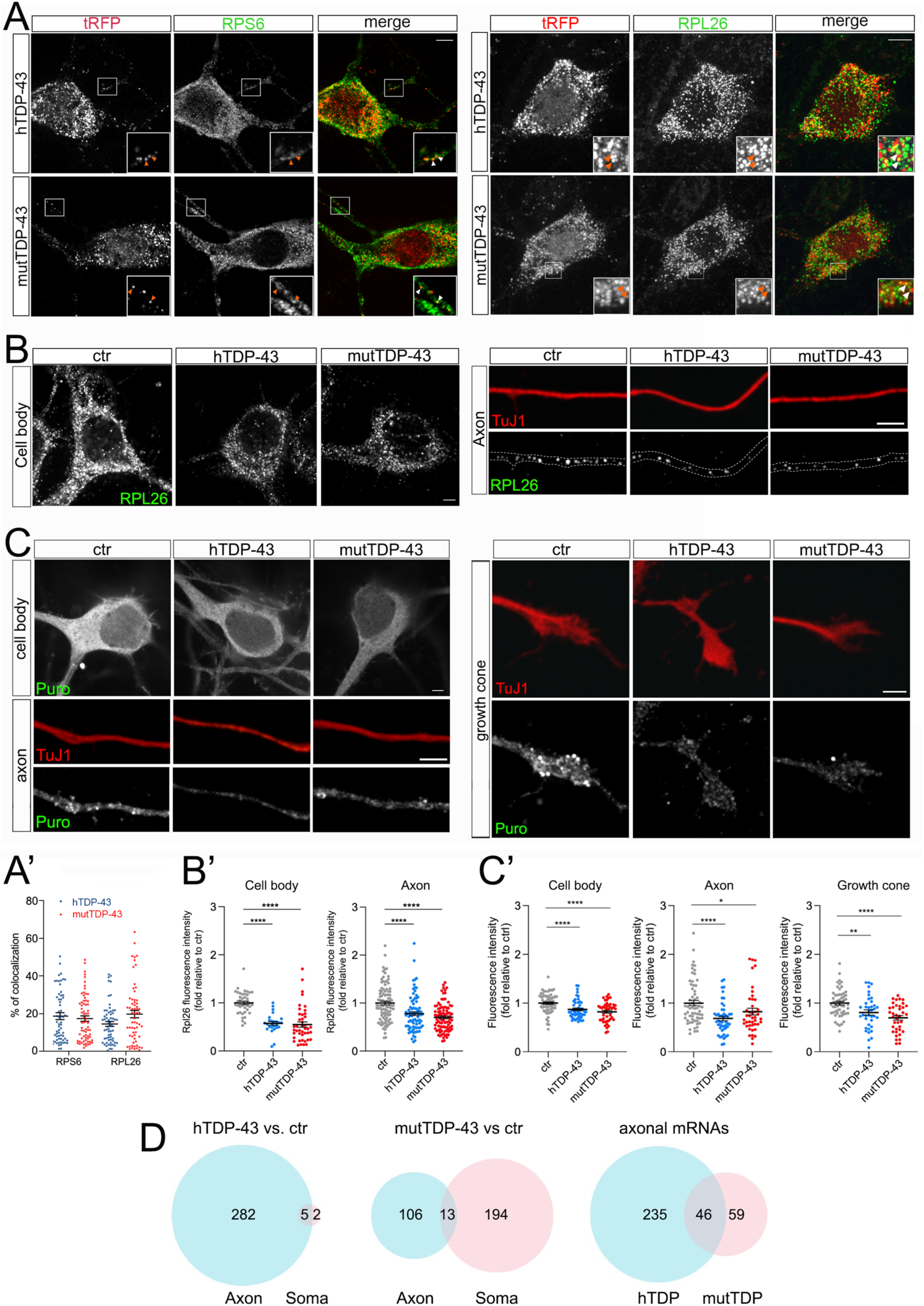
Global decrease of axonal protein synthesis in neurons expressing hTDP-43 and mutTDP-43. (A) A fraction of small and large subunit ribosomal proteins (RPS6 and RPL26, respectively) colocalize with aggregates of RFP-tagged human TDP-43, both wild type and mutant (see arrowheads in insets). (A’) dots in histograms indicate the percentage of colocalization in each individual cell examined (Mean±SEM, n=3, two-way ANOVA followed by Tukey’s test, n.s). (B) RPL26 protein abundance is reduced in the cell body (left) and axon (right) of cells expressing human TDP-43, wt or mutant. Differences and their statistical significance are plotted in B’, where each dot corresponds to an individual cell or axon examined. Mean ±SEM, n=3, Mann-Whitney test,****p-value<0.0001. (C) Global protein synthesis, expressed as the intensity of puromycin-positive signal, is decreased in the cell body, axon and growth cone of cells expressing human TDP-43, wt or mutant (see results for explanation). Differences and their statistical significance are plotted in C’, where each dot corresponds to an individual cell / axon / growth cone. (Mean±SEM, n=3, Mann-Whitney test, *p-value<0.05, **p-value<0.01, ****p-value<0.0001). (D) Here and in subsequent figures, the Venn diagrams on the left and in the center show the number of DEGs in the axon (light blue) and soma (pink) of hTDP-43 and mutTDP-43 neurons, respectively; the intersection between blue and pink diagrams on the right indicates axonal DEGs shared by the two populations, and listed in table 2. Venn diagrams shown here describe the numbers of polysome-engaged mRNAs, involved in various aspects of mRNA translation, whose abundance is decreased in hTDP-43 (wt) and mutTDP-43 (mut) axons vs ctr ones. Note the high number of axonal DEGs belonging to gene ontologies related to mRNA translation (see Methods).

Therefore, we set out to determine whether hTDP- and mutTDP-expressing neurons showed decreased axonal levels of RPL26, a component of the ribosomal large subunit for which efficient antibodies are available, relative to control axons. To this end, we immunostained hTDP-43, mutTDP-43 and ctrl neurons, cultured in microfluidic chambers for 9 DIV, with antibodies to RPL26 and the axonal marker TuJ1. RPL26 signal was then quantified in the axon and cell body. As shown in **Fig. 4B,B’**, RPL26 protein levels are decreased in both axons and cell bodies of hTDP-43 and mutTDP-43 neurons relative to ctr neurons. In conclusion, the decrease of axonal RPL26 suggests that the overall amount of 60S ribosomal subunit may be significantly decreased in hTDP-43 and mutTDP-43 axons and cell bodies, relative to ctr neurons.

Accordingly, we asked if protein synthesis might be globally defective in these neurons. To assess whether mRNA translation is affected in hTDP-43 and mutTDP-43 neurons, we carried out a puromycylation assay (David et al., 2012). By quantitative immunofluorescence methods, we evaluated puromycin incorporation as a measure of ongoing protein synthesis. As shown in **Fig. 4C,C’**, in both hTDP-43 and mutTDP-43 neurons puromycin signal is significantly reduced in the cell body, axon and growth cone alike, similar to findings previously observed at the neuromuscular junction (Altman et al., 2021).

To identify transcripts downregulated in the axon of hTDP-43 and mut-TDP-43 neurons, we performed next-generation sequencing (NGS) of the axon- and cell-body-specific polysome-engaged mRNAs (polysome profiling) from TDP-43-overexpressing neurons was performed by tag-free polysome isolation through a miniaturized sucrose gradient (see Methods, Negro et al., 2018). This approach consists of the following steps: i) hTDP-43, mutTDP-43 and ctr neurons are plated in microfluidic chambers for 9 DIV; ii) axonal and cell body compartments of these neurons are lysed and loaded onto a miniaturized sucrose gradient to separate polysome-engaged mRNAs from sub-polysomal mRNAs; iii) polysomal mRNAs and sub-polysomal mRNA fractions are isolated; iv) mRNAs are extracted from these fractions and sequenced (**Supplemental Fig. 4**).

We found that polysomal mRNA levels are robustly dysregulated in both human TDP-43- expressing neuronal populations compared to control neurons, with 1598 and 1601 differentially expressed genes (DEGs) for hTDP-43 and mutTDP-43, respectively. Compared to the number of DEGs in the cell body alone (467 in hTDP43 and 513 in mTDP43), more changes occur in the axon, accounting for 65% and 68% of total DEGs, respectively. Importantly, 1043/1131 (92%) axonal DEGs in hTDP-43 and 877/1088 (81%) in mutTDP-43 are downregulated, suggesting a widespread loss of mRNAs engaged in local translation (**Table 1**).

**Table 1.**

| TABLE1 |  | Cell body | Axon |
| --- | --- | --- | --- |
| **hTDP-43** | up-regulated | 86 | 88 |
|  | down-regulated | 381 | 1043 |
|  |  | **467** | **1131** |
| **mutTDP-43** | up-regulated | 85 | 211 |
|  | downregulated | 428 | 877 |
|  |  | **513** | **1088** |

Of all downregulated polysome-engaged transcripts, a large number belonged to gene ontologies related to protein synthesis and displayed lower levels in hTDP-43 and mutTDP-43 neurons (289 and 313, respectively), compared to control neurons (**Fig. 4D**). Of these, 47 axon-specific polysome-engaged DEGs were shared between the two overexpressing populations, and are listed in **Table 2**. Remarkably, in neurons overexpressing wt human TDP-43 (hTDP- 43), axon-specific polysomal DEGs make up 97.6% of the total, whereas in mutTDP-43 cells axon-specific DEGs amount 33.9 of the total.

**Table 2.**
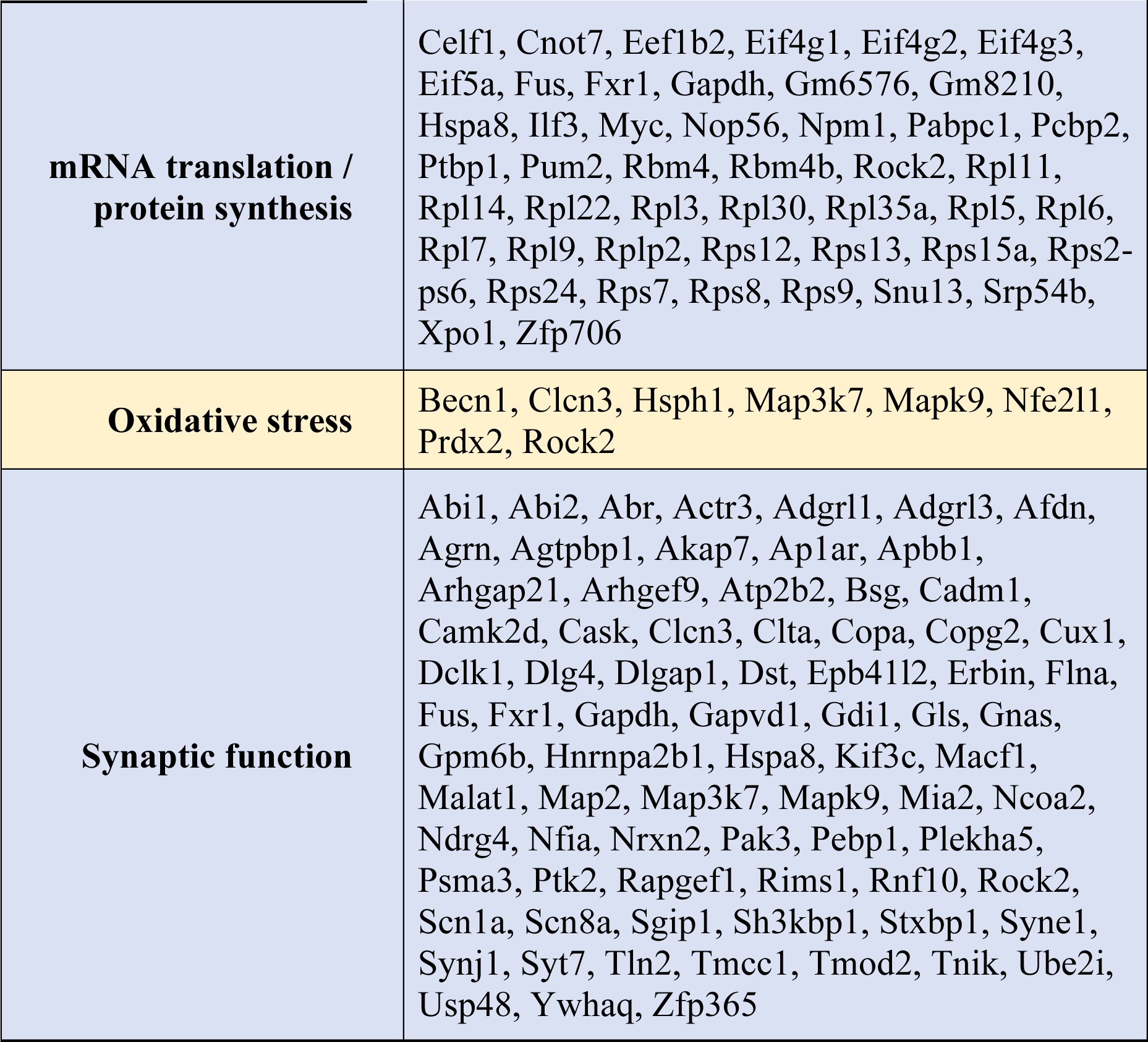
downregulated axonal mRNAs shared between hTDP-43 and mutTDP-43 populations.

### 3.5 Increased oxidative stress and apoptotic cell death in TDP-43-overexpressing neurons

Two other gene ontologies affected by TDP-43 overexpression were related to synaptic function and to the response to oxidative stress. The presence of high levels of ROS and the consequent oxidative stress-dependent neuronal damage are among the main hallmarks of ALS (Barber and Shaw, 2010). Therefore, we asked whether our neuronal model of TDP-43 proteinopathy exhibits this characteristic aspect of neurotoxicity. We performed this analysis in neurons loaded with H_2_DCFDA, both under untreated conditions (**Fig. 5A**) and upon mild iron overload (48 hours in the presence of 20 μM ferric iron); the latter provides information about the ability of neurons to counteract and detoxify an oxidative condition, promoted here by the iron-catalyzed Fenton reaction. Not only did our results show a significant increase in basal ROS levels in both overexpressing populations compared to ctr neurons (**Fig. 5B**, red dot bars), but they also revealed a sharply increased oxidative effect of iron overload in TDP-43 overexpressing neurons versus ctr cells (**Fig. 5B**, green dot bars). As expected, in the presence of increased basal levels of oxidative stress, TDP-43-overexpressing cultures display a significantly increased frequency of apoptotic cells (**Supplemental Fig. 5**).

**Figure 5.**
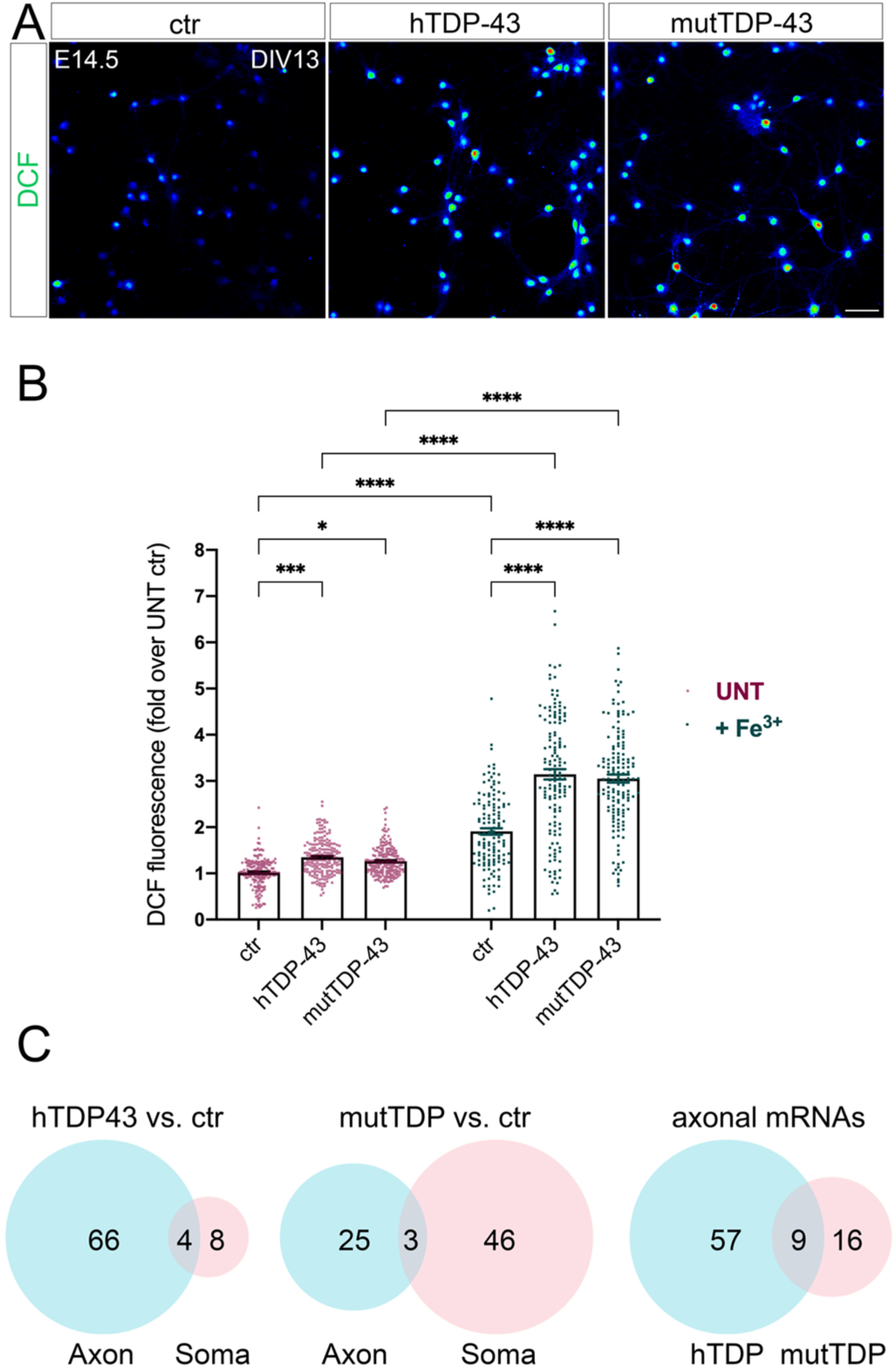
Impairment of the response to oxidative stress in neurons expressing human TDP-43, wt or mutant. (A) Representative images of ctr, hTDP-43 and mutTDP-43 neurons with incorporated DCF (green) shown as rainbow look-up table (LUT) values. (B) Graph showing DCF fluorescence intensity of hTDP-43, mutTDP-43 relative to ctr neurons of both untreated (UNT) and iron-treated (+Fe^3+^) cells. Data are expressed as mean ±SEM, from 4 or 5 biological replicates; each dot corresponds to an individual cell; Mann Whitney U-test, *, p<0.05; ***, p<0.005; ****p<0.0001. (C) Venn diagrams reporting the numbers of polysome-engaged mRNAs involved in the response to oxidative stress whose abundance is decreased in hTDP-43 (wt) and mutTDP-43 (mut) axons vs ctr ones. Note the high number of mRNAs, belonging to gene ontologies related to oxidative stress, which show differential abundance in the axon (see Methods).

As mentioned, numerous polysome-engaged somatic and axonal mRNAs involved in the response to oxidative stress are downregulated in neurons expressing hTDP-43 (78) and mutTDP-43 (74), compared to ctr neurons. Notably, a majority of differences observed in hTDP-43 neurons were relative to axonal mRNAs, suggesting that the axon plays a major and as yet uncharacterized role in the detoxification of oxidative stress. Out of the axonal DEGs, 9 were shared between hTDP-43 and mutTDP-43 neurons (**Fig. 5C** and **Table 2**), and include transcripts encoding nucleotide exchange factors for chaperone proteins, kinases, ER membrane sensors involved in stress response and *Prdx2*. This transcript encodes an abundant neuronal peroxidase that plays a key role in detoxifying hydrogen peroxide and therefore protecting neurons from oxidative stress (Bettegazzi et al., 2019); its reduced level in ALS neuronal axons might therefore contribute to functional alterations of UMNs (Liu et al., 2020).

### 3.6 Impaired spontaneous electrical activity, calcium handling and synaptic function in TDP-43 overexpressing neurons

As mentioned, gene ontologies related to synaptic function are affected in hTDP-43 and mutTDP-43 neurons, compared to control neurons. Synaptic abnormalities have been reported in the neurons of both ALS patients and animal models (primarily *SOD1* transgenics), ranging from morphological alterations to functional impairment, thereby supporting the hypothesis that ALS might either result from, or be exacerbated by synaptic dysfunction (Fogarty, 2019).

Since synaptic alterations seem to involve primarily glutamatergic neurons, as suggested by recent genetic studies (van Rheenen et al., 2021), we investigated whether hTDP-43 and mutTDP-43 overexpression could induce synaptic impairment in our cultures. Whole-cell patch clamp experiments revealed spontaneous, AMPA-dependent mini excitatory postsynaptic currents (mEPSCs), recorded at a holding potential of −70 mV and in the presence of a Na^+^ channel blocker (TTX, 1 μM) and GABA_A_ receptor antagonist (gabazine, 10 μM). No significant differences were detected in the average amplitude, decay time constant, or frequency of mEPSCs (**Supplemental Fig. 6**). However, spontaneous synaptic activity in TTX- and gabazine-free ACSF revealed recurrent bursts of postsynaptic currents (**Fig. 6A**). Interestingly, while burst duration was not affected by TDP-43 overexpression (either wild-type or mutant), the frequency of bursts was significantly reduced in both overexpressing populations compared to ctr neurons (**Fig. 6A**).

**Figure 6.**
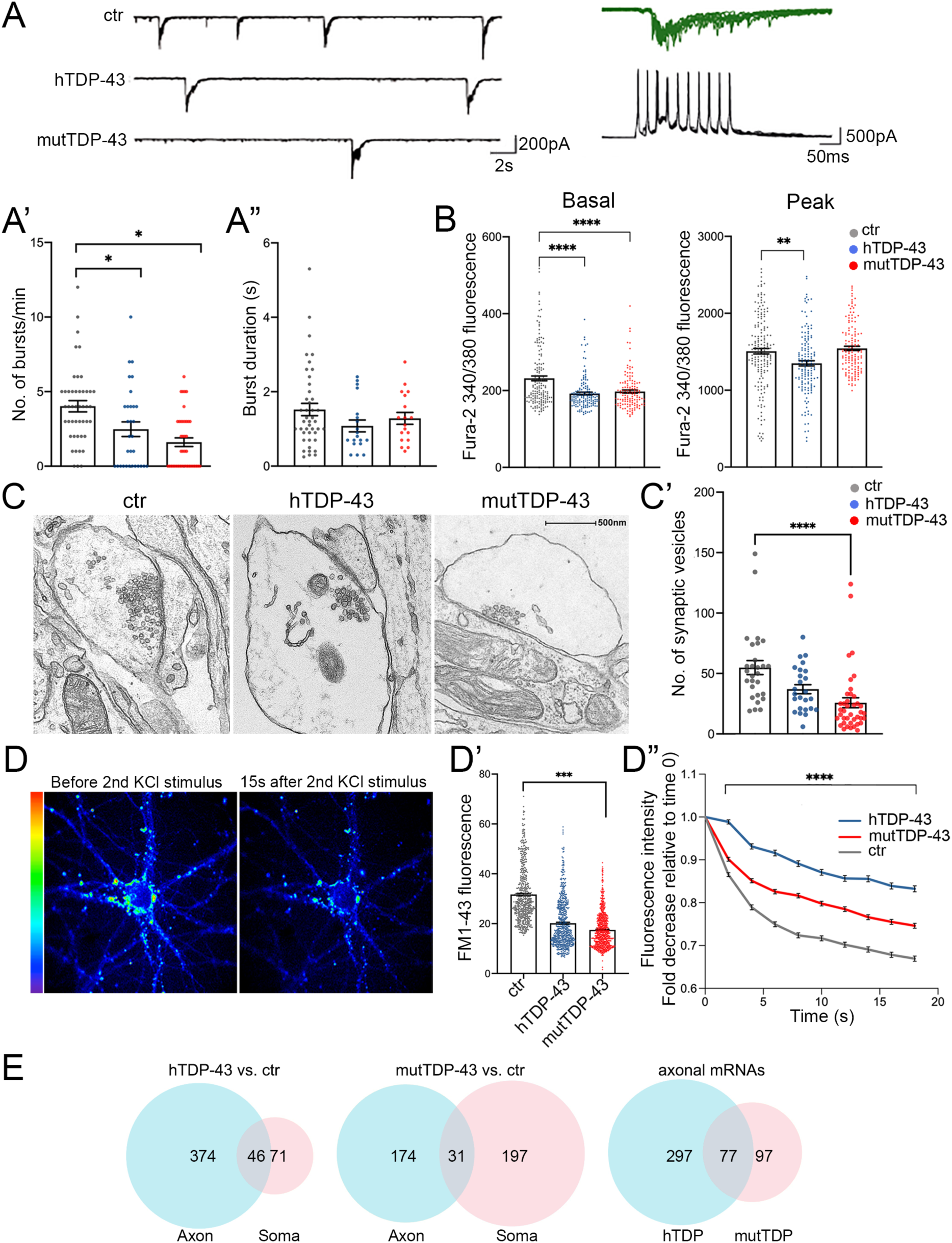
Impairment of spontaneous burst firing, calcium responses and synaptic vesicle exocytosis in neurons expressing human TDP-43. (A) Top left: examples of spontaneous compound postsynaptic current (PSC) bursts recorded in ctr, hTDP-43, and mutTDP-43 cultures. Top right: individual PSC bursts recorded in voltage clamp mode at a Vh of −70 mV correlate with spontaneous action potential bursts recorded in current clamp in a different cell. (A’) Summary plots of average burst frequencies. (A”) Average duration of individual bursts. hTDP-43 and mutTDP-43 show decreased burst frequency relative to ctr, while no difference in individual burst duration was observed. (Mean ±SEM, from 6 biological replicates, Mann Whitney U-Test, ns=p>0.05, *p<0.05). (B) Graphs showing fura-2 340/380 nm ratio of fluorescence intensity in ctr, hTDP-43 and mutTDP-43 neurons under basal conditions (graph on the left) and at the peak of response after glutamate stimulation (graph on the right); hTDP-43 and mutTDP-43 neurons show decreased calcium levels under basal conditions and at the peak of the response to glutamate stimulation (hTDP-43 only). (Mean ±SEM of 485 cells, from 5 biological replicates, Mann-Whitney test,**p<0.001, ****p<0.0001). (C) Transmission electron microscopy analysis performed on ctr, hTDP-43 and mutTDP-43 neurons. Images display synaptic boutons and the corresponding postsynaptic side. Note a significant reduction of vesicle numbers in mutTDP-43 neurons (C). (D) On the left, representative images showing FM1-43 fluorescence intensity as rainbow LUTs at two time points of the FM1-43 assay. The first image shows a neuron after treatment with 10 μM FM1-43 and 30 mM KCl simultaneously for 2 min to stimulate endocytosis and promote internalization of the dye in the intracellular vesicles, followed by a series of washes. The second image shows FM1-43 fluorescence intensity 15s after the second stimulus with 30 mM KCl, which triggers exocytosis of fluorescence-loaded vesicles. On the left, graph showing the decay of FM1-43 fluorescence relative to time 0, upon addition of the 2nd KCl stimulus, in hTDP-43 (blue), mutTDP-43 (red) and ctr cells (gray). (D’) the graph shows fluorescence levels of synaptic contacts upon completion of endocytosis (e.g. left panel in D), revealing a trend of FM1-43 signal decrease in hTDP-43 neurons and a significant decrease in mutTDP-43 cells. This result parallels the number of vesicles loaded with FM1-43. (D“): The downward slope of the curve parallels the rate of exocytosis, which is slower in TDP-43 overexpressing neurons (Mean ±SEM, from 5 biological replicates, Mann-Whitney test, ****p-value<0.0001, number of analyzed ROI: ctrl=456, wtTDP=384, mutTDP=532). (E) Venn diagrams reporting the numbers of polysome-engaged mRNAs involved in synaptic functions whose abundance is decreased in hTDP-43 (wt) and mutTDP-43 (mut) axons vs. ctr ones. Note the high number of differentially abundant mRNAs, belonging to gene ontologies related to synapses, which show differential abundance in the axon (see Methods).

The decrease of spontaneous bursting activity observed in TDP-43 overexpressing neurons prompted us to investigate the role of calcium (Ca^2+^) homeostasis in UMNs. To this end, the neurons were loaded with the fura-2 calcium dye which, thanks to its ratiometric properties, makes it possible to compare intracellular Ca^2+^ levels in overexpressing vs. control neurons (see Methods), both at rest and at maximal glutamate (100 μM) stimulation. In keeping with the decreased burst frequency, both hTDP-43 and mutTDP-43 UMNs showed decreased basal levels of intracellular Ca^2+^ concentration [Ca^2+^]_i_ compared to ctr neurons, therefore potentially affecting spontaneous electrical activity. Notably, the amplitude of [Ca^2+^]_i_ response to glutamate stimulation was also significantly reduced in hTDP-43 but not in mutTDP-43 neurons (**Fig. 6B**).

The impairment of spontaneous synaptic activity correlates with a depletion of vesicles in presynaptic active zones in TDP-43-overexpressing neurons, as revealed by ultrastructural analysis (**Fig. 6C**) and by the quantification of presynaptic vesicle numbers (graph in **6C’**). To confirm in live neurons the results obtained by electron microscopy, we exploited the properties of the FM1-43 dye (Amaral et al., 2011), since the loading and loss of the fluorescent dye provide a reliable measurement of synaptic vesicle retrieval and release in cultured neurons (Sambri et al., 2020). Exo-endocytosis was evoked by a first pulse of depolarizing stimulus (30 mM KCl added to the bath together with FM1-43 dye; see Methods); after washing the non-vesicle bound FM1-43, we quantified the fluorescence signal within synaptic boutons, correlating with their content of endocytic FM1-43-labeled vesicles (see image in **Fig. 6D**, left image). The graph in **Fig. 6D’** shows a trend of signal decrease in hTDP-43 neurons and a sharp decrease in mutTDP-43 cells, indicating, as in graph **6C’,** a lower number of vesicles in TDP-43-overexpressing neurons compared to ctr ones. The FM1-43-based experiments also allowed us to analyze the rate of synaptic exocytosis; indeed, after inducing fluorescent dye endocytosis, a second depolarizing KCl pulse caused vesicle exocytosis, evaluated by FM1-43 fluorescence decay over 20 seconds. As shown in **Fig. 6D** (right) **and 6D”**, mutTDP-43 and, even more so, hTDP-43 neurons, where calcium alterations are more pronounced, show a major impairment in their rate of exocytosis compared to ctr cells.

Again, a high number of DEGs was observed in the axonal translatome, underlying the key contribution of axonal protein synthesis in the regulation of synaptic activity and maintenance (**Fig. 6E**). The list of significantly downregulated synaptic mRNAs shared by hTDP-43- and mutTDP-43 axons includes transcripts encoding synaptic adhesion molecules, regulators of cytoskeletal dynamics, regulatory and structural proteins involved in synaptic vesicle trafficking/recycling; remarkably, some of the axon-specific differentially abundant mRNAs are also active in dendrites and postsynaptic terminals (**Table 2**).

## 4. DISCUSSION

In this paper, we combined the generation and functional characterization of mouse cortical neurons, used as cellular models of TDP-43 proteinopathy, with an analysis of their axonal translatome.

TDP-43 proteinopathy, manifesting itself as nuclear depletion and cytoplasmic accumulation / aggregation of TDP-43, is a very frequently observed feature of ALS (Arai et al., 2006; de Boer et al., 2020; Neumann et al., 2006). While animal models displaying physiological levels of TDP-43 aim at faithfully reproducing the pathophysiology of the disease, they often show a mild phenotype that can be properly analyzed only at late stages. The Q331K TDP-43 knock-in mouse model (White et al., 2018) shows almost no motor dysfunction nor TDP-43 proteinopathy, although it exhibits a cognitive phenotype at 12 months of age. Likewise, a knock-in model, carrying the M323K mutation, exhibits a mild muscular phenotype with reduced grip strength at 2 years of age and no TDP-43 pathology in the spinal cord or brain; however, at the molecular level, these mice show alterations in RNA splicing (Fratta et al., 2018). In this paper we describe and characterize primary neuronal models exhibiting spontaneous cytoplasmic TDP-43 aggregate formation, recapitulating, within a few days in culture and in a highly reproducible fashion, some cellular and functional aspects of ALS neuropathology that take decades to appear in patients. In our view, such models provide a solid screening platform and produce new evidence amenable to validation in more physiological model systems.

Primary cortical neuron cultures may contain a heterogeneous neuroglial population, which makes it difficult to discriminate neuronal cell-autonomous mechanisms from the effects of interactions between different cell types. Importantly, the neuronal cultures described here are highly homogeneous and strongly enriched in TBR1-positive glutamatergic projection neurons of the deep cortical layers, the cerebral neuron type that is most sensitive to ALS and FTD (Limone et al., 2021), in the virtual absence of GABAergic neurons or glial cells. In this model, exogenous tRFP-TDP-43 mainly localizes to the cytoplasm, generating mRNA granules, reminiscent of the pathological aggregates observed in the spinal cord of familial and sporadic ALS patients (Diaz-Garcia et al., 2021). However, while HuC/D is a hallmark and an established component of RNA granules and polysomes, defining the complete nucleotide and peptide composition of those aggregates will require further studies. In fact, mRNA granules recruit multiple factors, including RNA-binding proteins (Sidibé and Vande Velde, 2019).

Although the use of fluorescently tagged TDP-43 has been reported to impair its nuclear egress (Ederle et al., 2018), this is not the case in our models. Instead, we hypothesize that the size of the RFP-TDP-43 fusion proteins may reduce their nuclear import, causing trafficking defects akin to those produced by NLS removal (Igaz et al., 2011), and may cause a greater propensity to form cytoplasmic TDP-43 aggregates, unlike RFP alone. This feature allows early and stable formation of cytoplasmic aggregates that faithfully mimic TDP-43 proteinopathy. However, a quota of our fusion protein does localize to the nucleus, and negatively modulates the levels of the endogenous mouse TDP-43, likely due to the well established negative regulation exerted by TDP-43 on *TARDBP* mRNA stability (Avendano-Vazquez et al., 2012; Ayala et al., 2011).

Our results suggest that RFP-TDP-43+ granules sequester protein complexes required for mRNA translation, thus reducing the global rate of protein synthesis in the cell body, dendrites and axon alike. In keeping with this observation, RPL26 protein levels are profoundly reduced in the axon of hTDP-43 and mutTDP-43 neurons. Moreover, the results of our puromycylation assay directly show an overall decrease in axonal protein synthesis; in fact, the levels of numerous polysome-engaged mRNAs involved in protein synthesis are significantly decreased in the axon of TDP-43 overexpressing neurons. Such mRNAs are deregulated in the axons of both overexpressing populations and encode, among others, translation initiation factors and RNA-binding proteins. One open question regards the role of transcripts encoding ribosomal proteins in the axon (Nagano et al., 2020); recent evidence suggests that they may subserve the repair of damaged ribosomes (Fusco et al., 2021).

Our functional analysis also revealed that hTDP-43 and mutTDP-43 neurons exhibit increased oxidative stress levels, a hallmark of ALS neurons (reviewed in Barber and Shaw, 2010), and they do so in the absence of glial cells, suggesting that a neuronal-autonomous/paracrine mechanism contributes to the alteration of redox homeostasis in neurons displaying TDP-43 proteinopathy. Importantly, hTDP-43 and mutTDP-43 neurons show a basal increase of ROS production, indicating a chronic oxidative stress condition; moreover, these neurons display a reduced ability to counteract a condition of iron overload, used here to mimic the signs of iron accumulation that have been detected in the motor cortex (Bhattarai et al., 2020), frontal operculum, and precentral gyrus (Mitani et al., 2021) of ALS patients. Iron overload may, in turn, affect not only neuronal reducing potential, but also calcium homeostasis and synaptic function (Pelizzoni et al., 2008). We found that several mRNAs relative to oxidative stress response are underrepresented in the axon of hTDP-43 and mutTDP-43 neurons - in fact the number of downregulated axonal mRNAs exceeds that of deregulated transcripts in the soma of hTDP-43 neurons. It is reasonable to hypothesize that axonal translation may contribute importantly to the control of oxidative stress and that, in tract neurons projecting very long axons, this contribution may be extremely important.

Among the downregulated axonal mRNAs, *Prdx2* is highly expressed in normal spinal motoneurons (Kato et al., 2004) and plays an essential role in detoxifying hydrogen peroxide. Indeed, in ALS patients, *Prdx2* was found deregulated in surviving motoneurons at the intermediate disease stage, supporting its protective effect; on the other hand, the breakdown of this antioxidant system, occurring at late stages of ALS, accelerates neuronal death (Kato et al., 2005). This also occurs in the spinal cords of SOD1^G93A^ mice, where an increase of Prdx2 occurs during the onset of the disease, as a compensatory antioxidant defence, while its levels drop at late stage (Pharaoh et al., 2019).

With respect to synaptic activity, repeated bursts of postsynaptic currents, which represent a typical electrical activity pattern scored in mature cortical neuron cultures, were sharply and significantly decreased in our TDP-43 overexpressing populations, while burst duration only showed a nonsignificant decreasing trend. Burst activity is often mediated by an increase of intracellular calcium (Helmchen et al., 1999; Larkum et al., 1999; Williams and Stuart, 1999), and the decrease in burst frequency observed in hTDP-43 and mutTDP-43 neurons is in keeping with the finding that [Ca^2+^]_i_ is significantly reduced. This result is also in agreement with the deregulation of calcium ion homeostasis observed in different ALS models and proposed as a pathophysiological hallmark of the disease; indeed, in addition to an alteration of glutamate neurotransmission, several mechanisms involved in calcium handling (e.g. plasma membrane calcium extrusion, calcium influx through AMPA glutamate receptors, and calcium import from endoplasmic reticulum into mitochondria) have been found altered in ALS motor neurons (Guatteo et al., 2007; Sirabella et al., 2018). Moreover, a depletion of synaptic vesicles, revealed by EM in cultured neurons affects mutTDP-43 neurons more noticeably, contributing to the global impairment of synaptic functions of human TDP-43-expressing neurons.

In addition to the above, our results reveal that numerous polysome-engaged mRNAs encoding synaptic proteins are affected, and that their levels are altered both in the axon and cell body of human TDP-43-expressing neurons. Polysomal mRNAs deregulated in the axon of hTDP-43 and mutTDP-43 neurons control various aspects of presynaptic function, including cell adhesion, synapse formation, calcium responses, vesicular lumen acidification, glutamate synthesis, and neurotransmitter release; all these aspects will be addressed with complementary tools (Miesenbock et al., 1998).

While our functional analysis focused on exocytosis, the deficits observed in TDP-43-overexpressing neuronal populations may also affect endocytosis, as suggested by the reduced levels of transcripts involved in this stage of vesicle trafficking (Cremona et al., 1999; Mani et al., 2007; Miller et al., 2015).

Of note, a considerable number of downregulated transcripts involved in synaptic vesicle exocytosis (such as *Rab3a*, *Stx5a*, *Stx16*, *Snap25*, *Munc13*, *Rims1* and *Syt7*), are selectively downregulated in the axonal compartment of hTDP-43 neurons, in keeping with the observation that exocytosis is more severely impaired in neurons displaying cytoplasmic aggregates of wt TDP-43. However, the expression of mutant TDP-43 may cause a more global impairment, affecting neuronal maturation, supported by the significantly decreased number of synaptic vesicles scored in this population (Fig. 6C’).

More broadly, the results of our translatome analysis indicate that hTDP-43 expressing neurons display a selective impairment of axonal mRNA translation, whereas cells expressing mutTDP-43 exhibit a global mRNA deregulation, affecting the cell body and axon alike.

Finally, TDP-43+ granules recruit other RNA-binding proteins, such as FMRP, IMP and HuD (Fallini et al., 2012), while FUS, for instance, controls the splicing of a largely nonoverlapping set of transcripts (Lagier-Tourenne et al., 2012). Thus, the findings described here leave a key question unanswered, namely whether translation downregulation is restricted to TDP-43-bound mRNAs, possibly increasing the overall translation capacity for TDP-43-unbound transcripts.

## 5. CONCLUSIONS

This paper presents the results of multiple functional analyses performed on two cellular models of TDP-43 proteinopathy, the commonest neuropathological counterpart of ALS. The changes observed in our study correlate with the deregulation of axonal translation, further supporting its importance in the context of numerous homeostatic and functional processes. In particular, the prompt availability of proteins that are localized to the axon via slow axonal transport (reviewed in Roy, 2020) may be critically dependent on local mRNA translation. This, among other factors, may account for the high sensitivity of long-range motor neurons to ALS.

## Supporting information

Supplemental Figures Pisciottani et al 2023

## ACKNOWLEDGEMENTS

We thank all members of the Consalez and Viero labs for critical discussion of our data. Thanks to Claudia Rivoletti for preliminary experiments. Alessandra Pisciottani was supported during her PhD studies thanks to a very generous donation by the Fronzaroli family. Image analysis was carried out at ALEMBIC, an advanced microscopy laboratory established by the San Raffaele Scientific Institute and University. We gratefully acknowledge the financial support of AriSLA (AxRibALS grant to GGC and GV) and Ministero della Salute (RF21-2766 to GGC).

## DATA AVAILABILITY

Data will be made available on request.

## AUTHOR CONTRIBUTIONS

AP generated lentiviral vectors and performed primary cortical neuronal cultures, WB, puromycylation assay and subcellular fractionation; she also contributed to electrophysiological analysis and calcium, ROS and FM1-43 measurements. LC generated lentiviral vectors and performed primary cortical neuronal cultures, WB, puromycylation assay and subcellular fractionation; she also supervised most of the experimental procedures. CM performed subcellular fractionation, WB and lentiviral preparation. ES and JMC performed colocalization analysis. ST performed electrophysiological studies. PP and AQ performed electron microscopy analysis. FL, MM and GV performed the transcriptome and translatome analyses. GV contributed importantly to writing of the manuscript. AA performed the statistical analysis. OC critically read the manuscript. FiC performed confocal microscopy analysis. FrC performed and supervised calcium, ROS and FM1-43 measurements, and wrote the manuscript. GGC conceived and designed the study, supervised the analyses, and wrote the manuscript.

All authors contributed to the article and approved the submitted version.

## Declaration of Competing Interest

The authors declare no competing interests.

## Notes

### Competing Interest Statement

The authors have declared no competing interest.

### Summary of Updates

Small changes of Figure 6, of the Figure 4 legend, and of the discussion

