## Supplemental Figures Pisciottani et al 2023 for "Two neuronal models of TDP-43 proteinopathy display reduced axonal translation, increased oxidative stress, and defective exocytosis"

Alessandra Pisciottani et al.

### SUPPLEMENTAL FIGURES

Supplemental figure 1

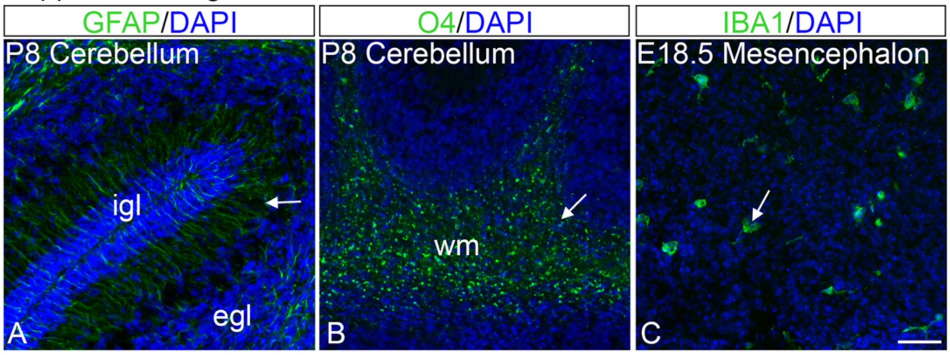


**Supplemental figure 1. Positive controls of glial markers.**
Immunofluorescence of cerebellar or mesencephalic sections as positive controls of the antibodies used on cell cultures (Fig. 1D). Sagittal sections of P8 mouse cerebellum show GFAP-positive signals (arrow) in Bergmann glia (A) and O4-positive cells (arrow) in white matter (B). A coronal section of E18.5 mouse mesencephalon shows IBA1-positive microglial cells (arrow) (C). igl: internal granular layer; egl: external granular layer; wm: white matter. Size bar: 50 μm.

Supplemental figure 2

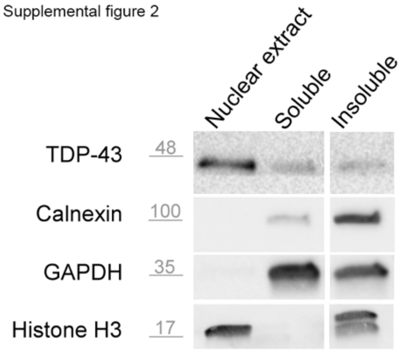


**Supplemental figure 2. Validation of the subcellular fractionation protocol (Fig. 2C).**

Western blot of the subcellular fractions of proteins extracted from non transduced cortical neurons at 14 DIV. Endogenous TDP-43 is mainly enriched in the nuclear fraction. Calnexin, a transmembrane protein of the endoplasmic reticulum, and the cytosolic protein GAPDH are enriched in the insoluble cytoplasmic fraction and in the soluble cytoplasmic fraction, respectively. Histone H3 is present in both nuclear and insoluble cytoplasmic fractions.

Supplemental figure 3

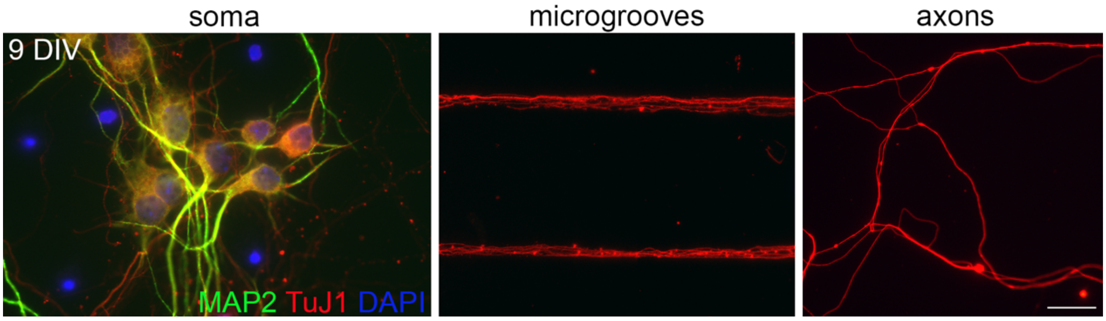


**Supplemental figure 3. Physical separation of whole-cell and axonal compartments in microfluidic chambers.**

Immunofluorescence of UMN cultures, grown in microfluidic chambers until 9 DIV, with antibodies for TuJ1 (red), a marker of dendrites and axons alike, MAP2 (green), a dendrite-specific marker, and DAPI, a nuclear marker. Microfluidic chambers allow the physical separation of axons (MAP2-negative) from dendrites (MAP2-positive) and cell bodies. Size bar: 20μm.

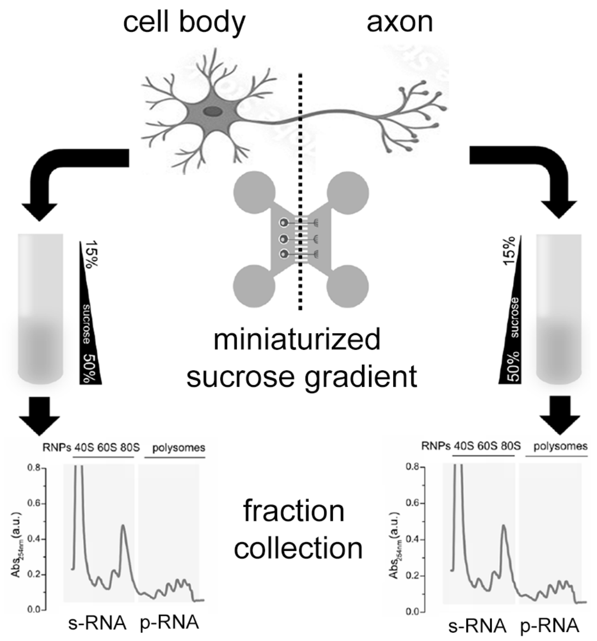


**Supplemental figure 4. Representative scheme of polysome-engaged and sub-polysomal mRNA isolation for sequencing.**

Neurons transduced with ctr, hTDP-43 and mutTDP-43 lentiviral particles were grown in microfluidic chambers until 9DIV. Axonal and whole-cell compartments were lysed and charged onto a miniaturized sucrose gradient. Then mRNAs from polysomal fractions (p-RNAs, containing polysome-engaged mRNAs) and subpolysomal fractions (s-RNAs, containing mRNAs not associated with polysomes) were isolated and sequenced.

Supplemental Figure 5

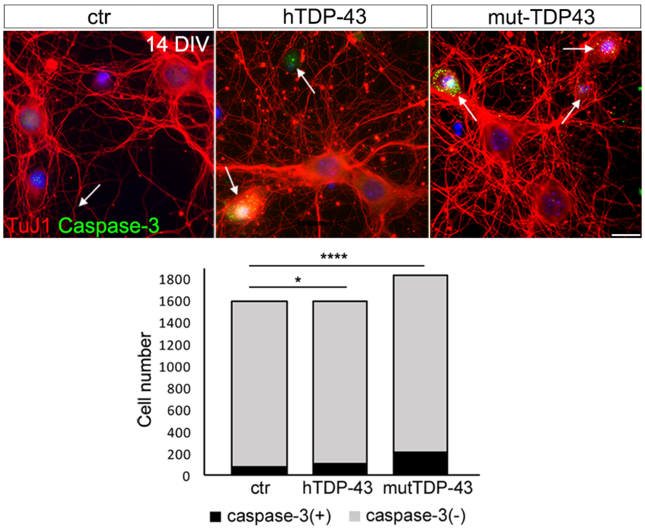


**Supplemental figure 5. hTDP-43 and mutTDP-43 show slight increased cell death.**

Immunofluorescence of ctr, hTDP-43 and mutTDP-43 (14 DIV), with anti-active Caspase 3 (in green) and anti-TuJ1 (in red) antibodies. The DAPI counterstaining (in blue) marks the cell nuclei. White arrows indicate active Caspase-3 positive cells. Size bar: 10µm. The histogram on the right shows the number of active Caspase-3 positive cells (in black) in hTDP-43, mutTDP-43 and control. The Caspase-3 negative cells are visualized in gray. The hTDP-43 and mutTDP-43 neurons show a statistically significant increase in the number of apoptotic cells. (n=5; Fisher's exact test; *$p<0.05$; ****$p<0.0001$).

Supplemental Figure 6

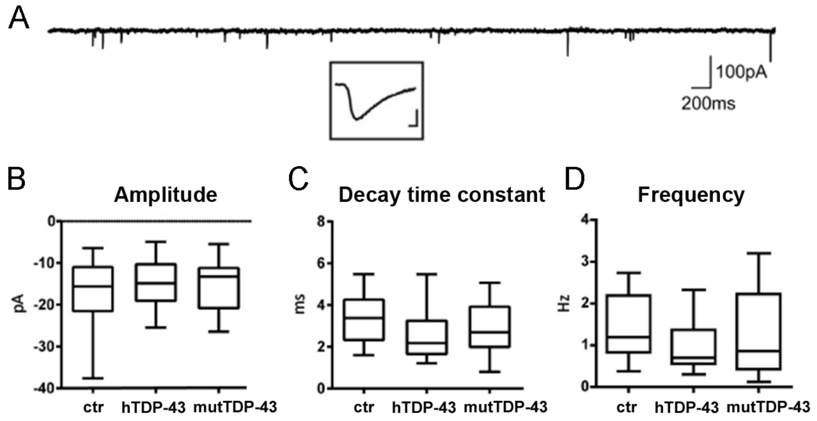


**Supplemental figure 6. hTDP-43 and mutTDP-43 neurons show no significant differences in mEPSC amplitude, decay time constant, and frequency.**
(A) Example of spontaneous mEPSCs recorded in voltage clamp mode (Vh: -70mV) in a cultured cortical cell. The inset shows an enlarged view of an individual mEPSC (scale bar: 10pA, 1ms). (B-D). Summary box plots of peak amplitude, decay time constant, and frequency of mEPSCs recorded in ctrl, hTDP-43, and mutTDP-43 cultures (box plots represent medians, 25 and 75 percentiles, and min-max values, from 6 biological replicates, Mann Whitney U-Test, ns=$p \geq 0.05$).

**Supplemental figure 7. Reduced vesicle number and impaired presynaptic organization in TDP-43-overexpressing neurons**
Transmission electron microscopy analysis performed on ctr, hTDP-43 and mutTDP-43. Images display synaptic boutons and the corresponding postsynaptic side. Note reduced vesicle number and inefficient clustering of presynaptic terminals in TDP-43-overexpressing neurons.
